# Rapid response to the Alpha-1 Adrenergic Agent Phenylephrine in the Perioperative Period is Impacted by Genomics and Ancestry

**DOI:** 10.1101/664961

**Authors:** Stephane Wenric, Janina M. Jeff, Thomas Joseph, Muh-Ching Yee, Gillian M. Belbin, Regeneron Genetics Center, CBIPM Genomics Group, Aniwaa Owusu Obeng, Stephen B. Ellis, Erwin P. Bottinger, Omri Gottesman, Matthew A. Levin, Eimear E. Kenny

## Abstract

**Background:** The emergence of genomic data in biobanks and health systems offers new ways to derive medically important phenotypes, including acute phenotypes that occur during in-patient clinical care. We hypothesized that there is a genetic underpinning to the magnitude of the response to phenylephrine, an α1-adrenergic receptor agonist commonly used to treat hypotension during anesthesia and surgery.

**Methods:** We quantified the response to phenylephrine by determining the delta between the minimum blood pressure (BP) within five minutes before and the maximum BP within five minutes after bolus administration. We then performed a genome-wide association study (GWAS) adjusted for genetic ancestry, demographics, and relevant clinical covariates to investigate genetic factors underlying individual differences systolic BP response to phenylephrine (ΔSBP), as well as mean arterial pressure (ΔMAP) and diastolic BP (ΔDBP), for both the entire study cohort as well as for each of 3 ancestry sub-cohorts; European American(EA), African American(AA), and Hispanic American(HA).

**Results:** 4,317 patients met inclusion criteria, of which 3,699 were genotyped. Average ΔBP values over the entire cohort were ΔSBP=17(+-25) mmHg, ΔMAP=14(+-18) mmHg, ΔDBP=11(+-14) mmHg. The largest difference between populations was observed for ΔSBP (ΔSBP_EA_=20(+-24) mmHg; ΔSBP_HA_=16(+-25) mmHg; ΔSBP_AA_=15(+-25) mmHg). The differences remained after adjusting for clinical covariates and ancestry (EA vs. HA: ΔSBP, p<0.032;ΔMAP, p<0.021;ΔDBP,p<0.008);(EA vs. AA:ΔSBP,p<5.13×10^-5^;ΔMAP,p<2.1×10^-4^;ΔDBP,p<3.3×10^-4^). GWAS revealed significant associations between loci and BP response in 5 different genome regions (p<5×10^-8^) in the entire cohort, and suggestive associations in 2 different regions in EAs (p<6×10^-8^,p<7×10^-8^). We observed non-random enrichment in association with SBP drug response in 165 loci previously reported to be associated with systolic blood pressure. Finally, we discovered rare variants, rs188427942 and rs147664194 present at ∼1% in EAs and rs146535276 present at ∼1% in AAs respectively, where patients carrying one copy of these variants show no response to phenylephrine.

**Conclusions:** It is possible to derive a quantitative phenotype suited for comparative statistics and genome-wide association studies from routinely collected perioperative data. There are population differences in rapid response to phenylephrine, large effect alleles and novel genes affecting pharmaceutical response, and phenylephrine non-responders, with implications for personalized treatment during surgery.

## Introduction

Perioperative phenotypes, such as rapid response to drugs administered during surgery, constitute an integral part of medical practice, as the variation in drug response across individuals is a fundamental variable to be taken into account before, during, and after surgery. One such perioperative phenotype is rapid physiologic responsiveness to short acting adrenergic agents administered to treat hypotension during surgery. Prolonged or recurrent episodes of hypotension during anesthesia and surgery are believed to be associated with increased morbidity and mortality (Bijker et al., 2009; Charlson et al., 1990; M. A. Levin et al., 2015; Monk, Saini, Weldon, & Sigl, 2005; Sessler et al., 2012; Vernooij et al., 2018; Walsh et al., 2013), thus the use of these adrenergic agents is common. In the case of response to adrenergic agents, although anecdotal evidence has led clinicians to be aware of individual variation in response between patients, the lack of a rigorous replicable and quantitative phenotype prevents population level comparisons as well as the use of tools from statistical genetics. We focused our investigation on phenylephrine, a selective α1-adrenergic receptor agonist, because it is one of the most commonly used agents for the treatment of intraoperative hypotension, with a very rapid onset when given intravenously and a short half-life of 15-20 minutes. In addition, there is a substantial body of work dating back to the early 1990’s investigating receptor variants (Amory et al., 2002; Flancbaum, Dick, Dasta, Sinha, & Choban, 1997; Kleine-Brueggeney, Gradinaru, Babaeva, Schwinn, & Oganesian, 2014; Oganesian, Yarov-Yarovoy, Parks, & Schwinn, 2011; Schwinn & Reves, 1989). We hypothesized that there would be a genetic basis that could help explain both the individual variability as well as the variation in response seen between individuals of differing ancestry.

## Methods

### Study Population

The Icahn School of Medicine at Mount Sinai (ISSMS) Department of Anesthesiology maintains a linked database called iGAS that combines data from our institutional Biobank (called Bio*Me*) and our electronic anesthesia record. The architecture and content of iGAS have been previously described. (Matthew A. Levin et al., 2017). The resulting linked database is a rich resource of high resolution, high quality perioperative data that can be used to generate robust phenotype profiles.

Procedures were categorized based on ICD-9 Procedure codes, using the Healthcare Cost and Utilization Project Clinical Classification Software (HCUP CCS, REF https://hcup-us.ahrq.gov/toolssoftware/ccs_svcsproc/ccssvcproc.jsp) to map individual procedures to high level procedure categories (e.g., Digestive System). The Charlson comorbidity index (CCI) was calculated for each patient using administrative ICD-9-CM discharge diagnosis codes and an ICD-9-CM to comorbidity map with revised diagnosis weights (Quan et al., 2005; Schneeweiss, Wang, Avorn, & Glynn, 2003). Calculation of the CCI was done using the R package *medicalrisk* (McCormick, 2015).

### Phenotypic modelling

Intraoperative blood pressure (BP) is typically measured either with a Non-Invasive Blood Pressure (NIBP) cuff at a frequency of once every 1-5 minutes, or continuously via invasive arterial line. Arterial line data are recorded as often as once every 15 seconds. Blood pressure measurements outside of normal physiological ranges, as determined by validity flags in the electronic anesthesia record, were excluded as they likely represented measurement artifacts. To derive a phenotype suitable for further analysis, we defined a rapid drug response as the *difference* in the recorded *minimum* blood pressure (systolic - SBP, diastolic - DBP, and mean - MAP) within a five-minute period *before* and the *maximum* blood pressure in the five-minute period *after* a bolus of phenylephrine (**Figure 1A**). Five-minute *before-* and *after-* windows were used to account for the fact that NIBP measurements and recordings are intermittent, as described above, and also that charting of bolus drug administration is often non-contemporaneous. For patients with an arterial line, this meant a measurement within as soon as 15 seconds before or after the drug dose was used. For patients with an NIBP, the closest reading could be as soon as within 1 minute, if the NIBP cycle time was set to its shortest value. In either case (NIBP or invasive pressure monitoring), the longest delay between a bolus dose and a BP reading would be a maximum of 5 minutes. This ensured that the time relationship between drug administration and BP measurement was reasonably constant across all patients. (**Figure 1)**. The final phenotypic measurement was thus the Δ SBP, DBP and MAP response to a bolus of phenylephrine, within 5 minutes of administration.

**Figure 1A.**
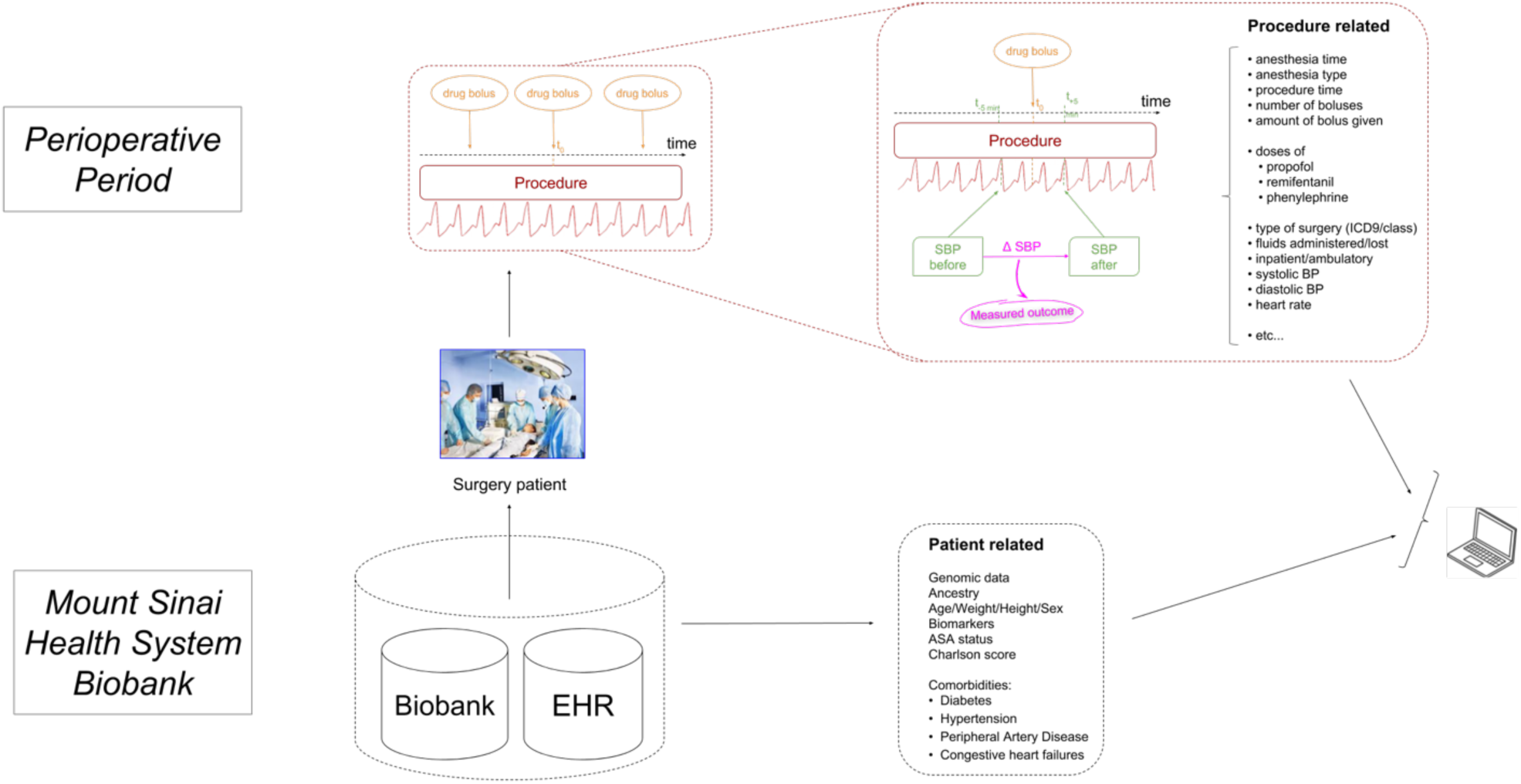
Data capture schema. The raw phenotype is captured during surgery within a 10-minute interval. Additional procedure-related information, as well as general patient-related variables, including genomic data, are combined through the analysis pipeline.

**Figure 1B.**
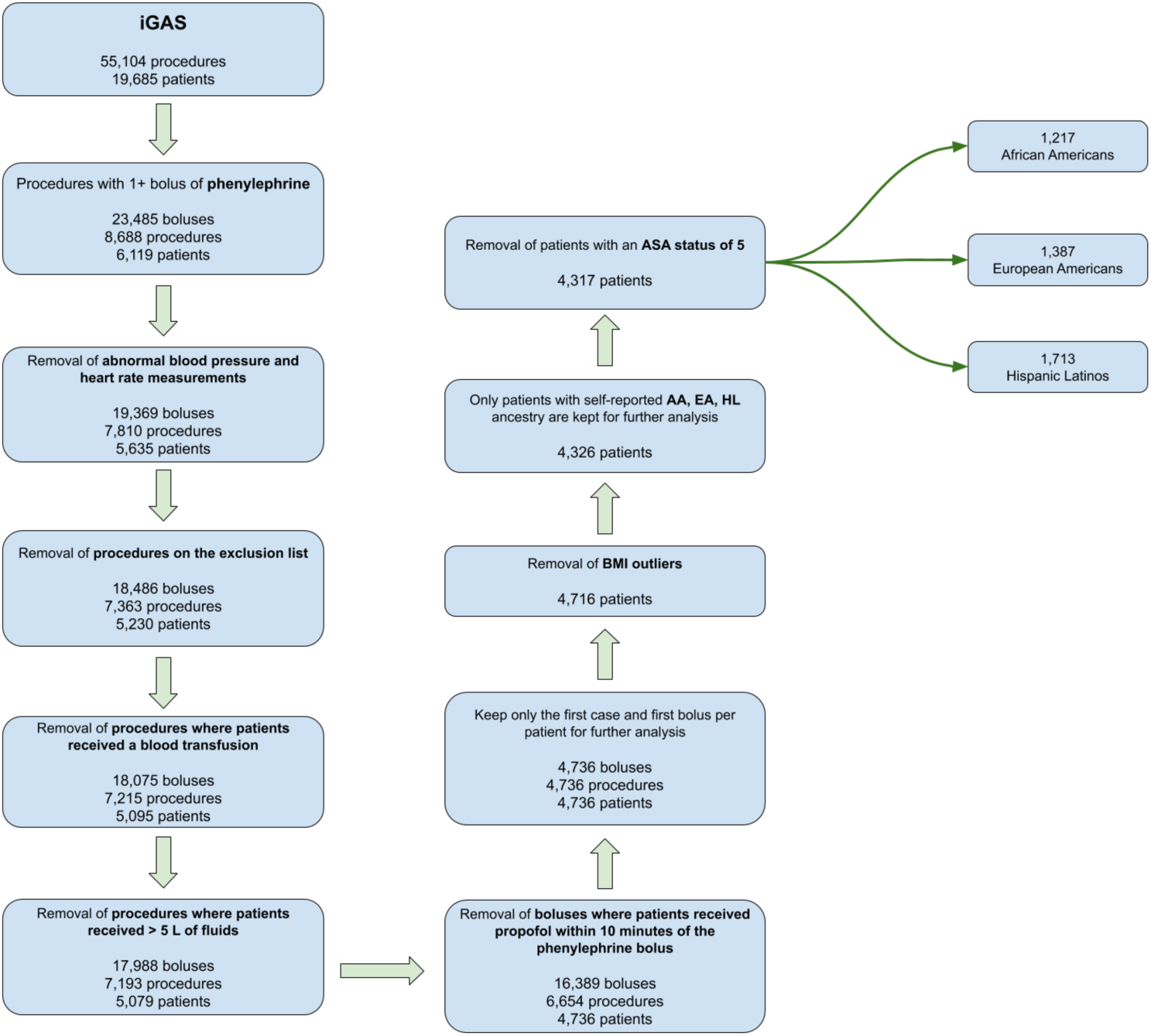
Inclusion/Exclusion criteria. From an initial cohort of 19,685 iGAS patients matched to 55,104 procedures, the sequential exclusion criteria and filtering steps reduced the number of patients included in the study to 4,317 comprising 1,217 African Americans, 1,387 European Americans and 1,713 Hispanic/Latinos.

### Case Inclusion/Exclusion criteria

Initial inclusion criteria were all Bio*Me* patients who had undergone anesthesia and received at least one bolus of phenylephrine intraoperatively. A diagram showing all exclusion steps is shown in **Figure 1B**. Cancelled cases or manually charted cases that had no automated recording of physiologic data were excluded. We also excluded all procedures of short duration and rapid turnover such as colonoscopies and endoscopies where drug bolus administration is known to often be significantly undercharted (Avidan, Dotan, Weissman, Cohen, & Levin, 2014; Wax & Feit, 2016). Emergency procedures (∼1%) were excluded, for several reasons: 1) these cases often involve blood loss and fluid shifts that may have profound effect on intravascular volume and responsiveness to phenylephrine, 2) emergency cases often receive blood products and fluid that are recorded with poor temporal resolution, and 3) charting of drug administration is often compromised and inaccurate during emergency situations. We also excluded procedures in which the patient received any blood transfusion, as blood is a potent vasoactive agent and intravascular volume expander, and the presence of blood transfusion was found to be highly correlated with phenylephrine drug response (**Supplementary Figure 1**, p-value = 5.40e-10). As the distribution of total crystalloid administered was skewed and found to be highly correlated with phenylephrine drug response (**Supplementary Figure 2**, p-value = 4.9e-06), we excluded procedures with > 5L of fluid administration charted (∼2%).

**Figure 2.**
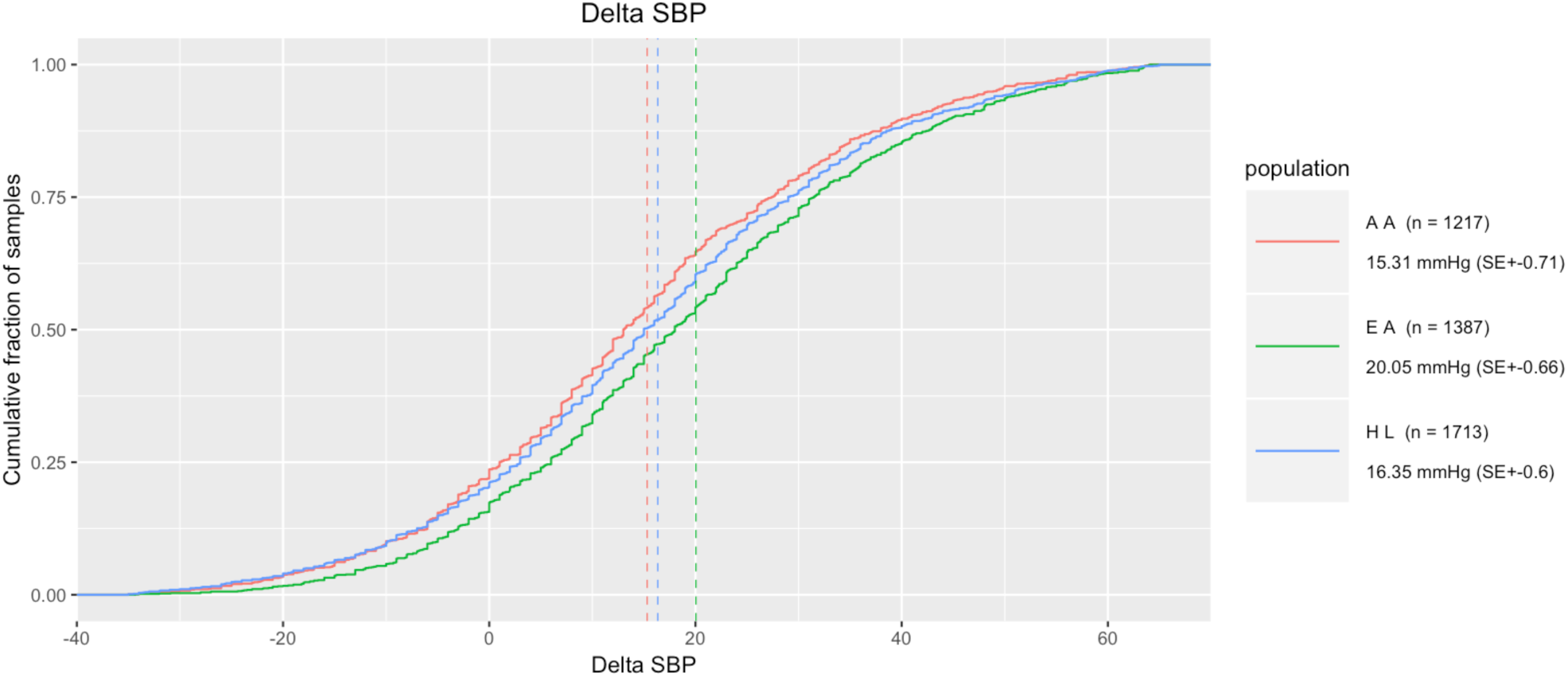
Drug response difference between populations. Cumulative distribution function for Δ Systolic Blood Pressure for African-Americans (in red), European Americans (in green) and Hispanic/Latinos (in blue).

Preliminary analysis demonstrated a large confounding effect of the intravenous anesthetic agent propofol on the phenylephrine signal (p-value = 5.764e-05 for a linear regression between Δ SBP and propofol bolus time). Propofol is known to cause a predictable and often significant decrease in blood pressure and has a similar onset (immediate) and duration (10 minutes) to that of phenylephrine. There was a significant difference in Δ SBP between patients with a propofol bolus administered within 10 min of a phenylephrine bolus and patients with no propofol bolus or propofol administered within more than 10 minutes of phenylephrine (Mann-Whitney p-value < 0.003). We therefore excluded phenylephrine bolus events within ten minutes of a bolus dose of propofol. If that phenylephrine bolus was the only bolus the patient received during their procedure, the entire procedure was excluded. For patients with more than one anesthetic/procedure in the data set, only data from the first anesthetic was used. In order to be able to study the inter-individual variability and to investigate the genetic basis of the phenotype, only one bolus-response was kept per patient. We kept the Δ SBP, DBP and MAP associated with the first case and first bolus per patient.

Finally, we excluded BMI outliers (BMI > 100, likely due to data entry error) and procedures on critically ill patients with an American Society of Anesthesiologists (ASA) Physical Status of 5 (not expected to survive without surgery, n = 9). Such patients usually present with severe organ system derangement and little to no sympathetic reserve. After all patients, procedures, and bolus-response exclusion steps, our cohort contained a total of 4,317 patients (1,217 AA, 1,387 EA, 1,713 HL), among which 3,699 were genotyped and passed QC metrics (1,082 AA, 1,112 EA, 1505 HL).

### Statistical Analysis

Univariate analysis was used to select covariates to include in the genome-wide association study (GWAS), with the level of significance set at *p-value = 0.05*. The distribution of total crystalloid administered remained correlated with phenylephrine drug response after exclusion of procedures with > 5L of fluid administered (**Supplementary Figure 2**, p-value = 4.9e-06). We observed a nominal association between BMI and drug response (p-value = 0.03), even after exclusion of outliers likely due to data entry errors. Age and gender were normally distributed in iGAS. Age was associated with drug response (p-value = 4.55e-14, **Supplementary Figure 3**) so age-adjusted adjusted z-scores were used for further analysis. Sex was only nominally associated with drug response (p-value = 0.07), thus no exclusions or adjustments were made. ASA status (p-value = 3.58e-07), depth of general anesthesia as measured by the Minimal Alveolar Concentration (MAC, p-value = 5.23e-10, **Supplementary Figure 4**) was associated with drug response in the univariate analysis. CCI was not associated with drug response. We found that phenylephrine response was slightly different between cardiovascular, musculoskeletal, and gastrointestinal surgeries, but the association was not statistically significant (p-value = 0.6125). The bolus amount of phenylephrine given was not associated with the phenotype; however, given the distribution of bolus amounts (**Supplementary Figure 5**), the likely non-linearity of the dose-response relationship, and the absence of per-patient dose-response curves, we kept phenylephrine bolus amount as a covariate in the model. To account for population level differences, a genetic relationship matrix was calculated and the first 10 principal components were included as covariates in the model (Conomos, Miller, & Thornton, 2015; Yang, Zaitlen, Goddard, Visscher, & Price, 2014). The final list of covariates used in the model for the association analysis were sex, BMI, ASA status, depth of anesthesia, phenylephrine bolus amount, total volume of fluid administered, and the first 10 principal components.

**Figure 3A.**
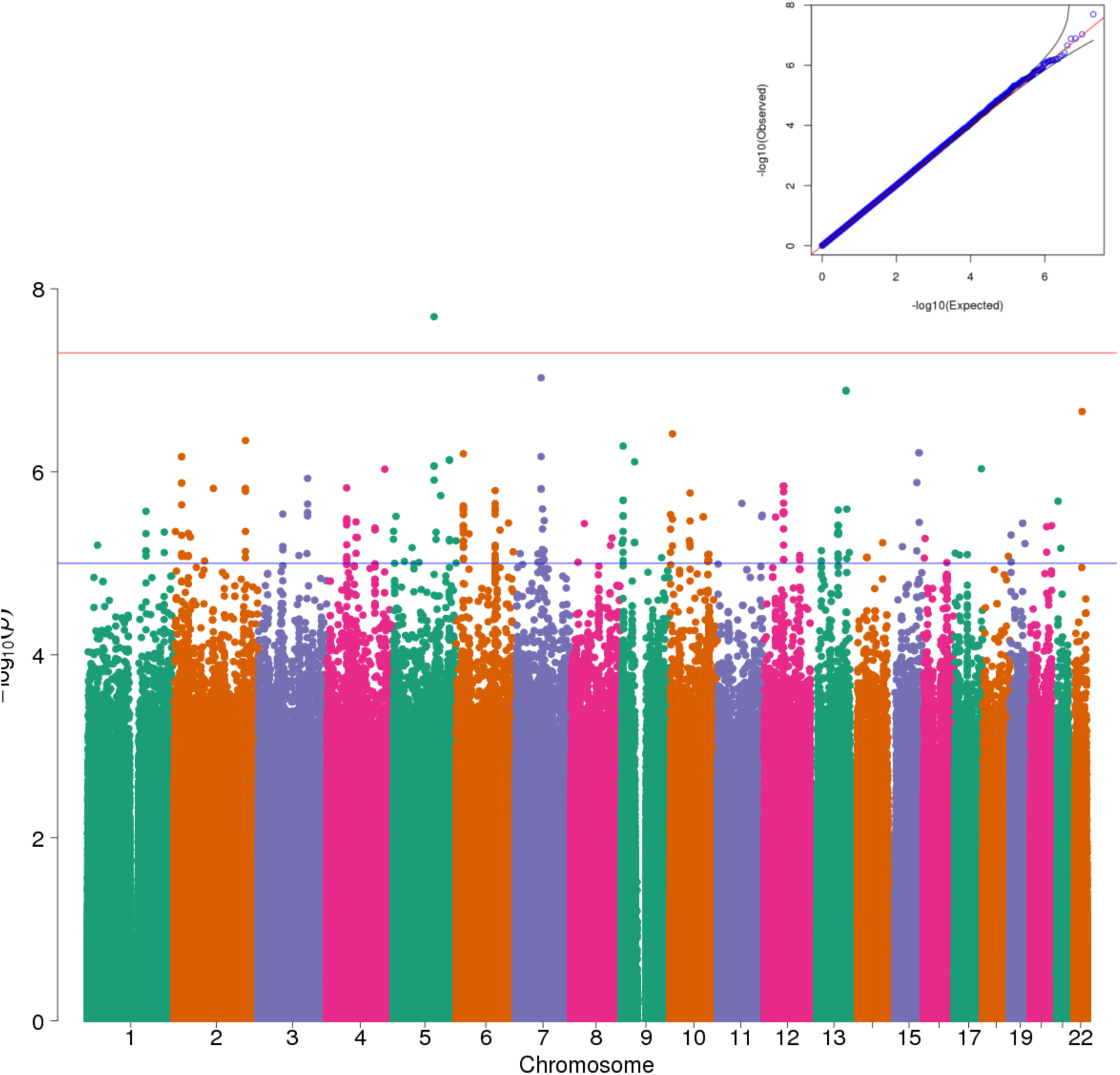
Manhattan plot and QQ plot of the genome-wide association study performed on Δ SBP with the whole cohort. The red horizontal line denotes the genome-wide significance threshold for p-values (5e-8) while the blue horizontal line denotes a p-value threshold of 1e-5. The top SNP is rs145337816, located on chromosome 5, at locus 117211218 (in hg19 coordinates). The minor allele frequency is 0.46%, the imputation info score is 0.909. The association p-value is 2.02e-8. Lambda-gc = 1.0059

**Figure 3B.**
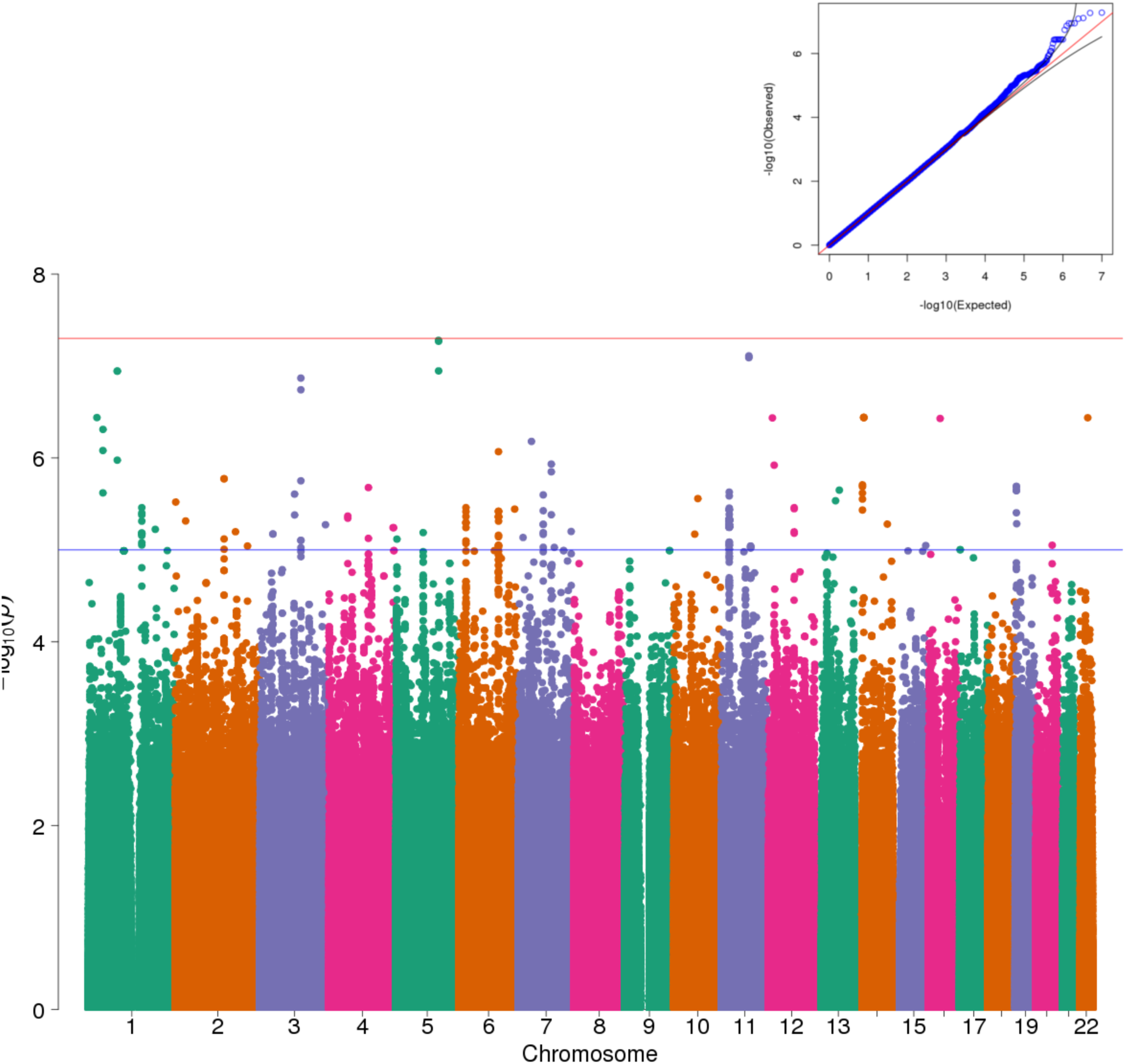
Manhattan plot and QQ plot of the genome-wide association study performed on Δ SBP with European Americans. The top SNP is rs188427942, located on chromosome 5, at locus 122900413 (in hg19 coordinates). The minor allele frequency is 1.16 % in EA (0.57% in the whole cohort), the imputation info score is 0.759. The association p-value is 5.26e-8. Lambda-gc = 0.9976.

**Figure 3C.**
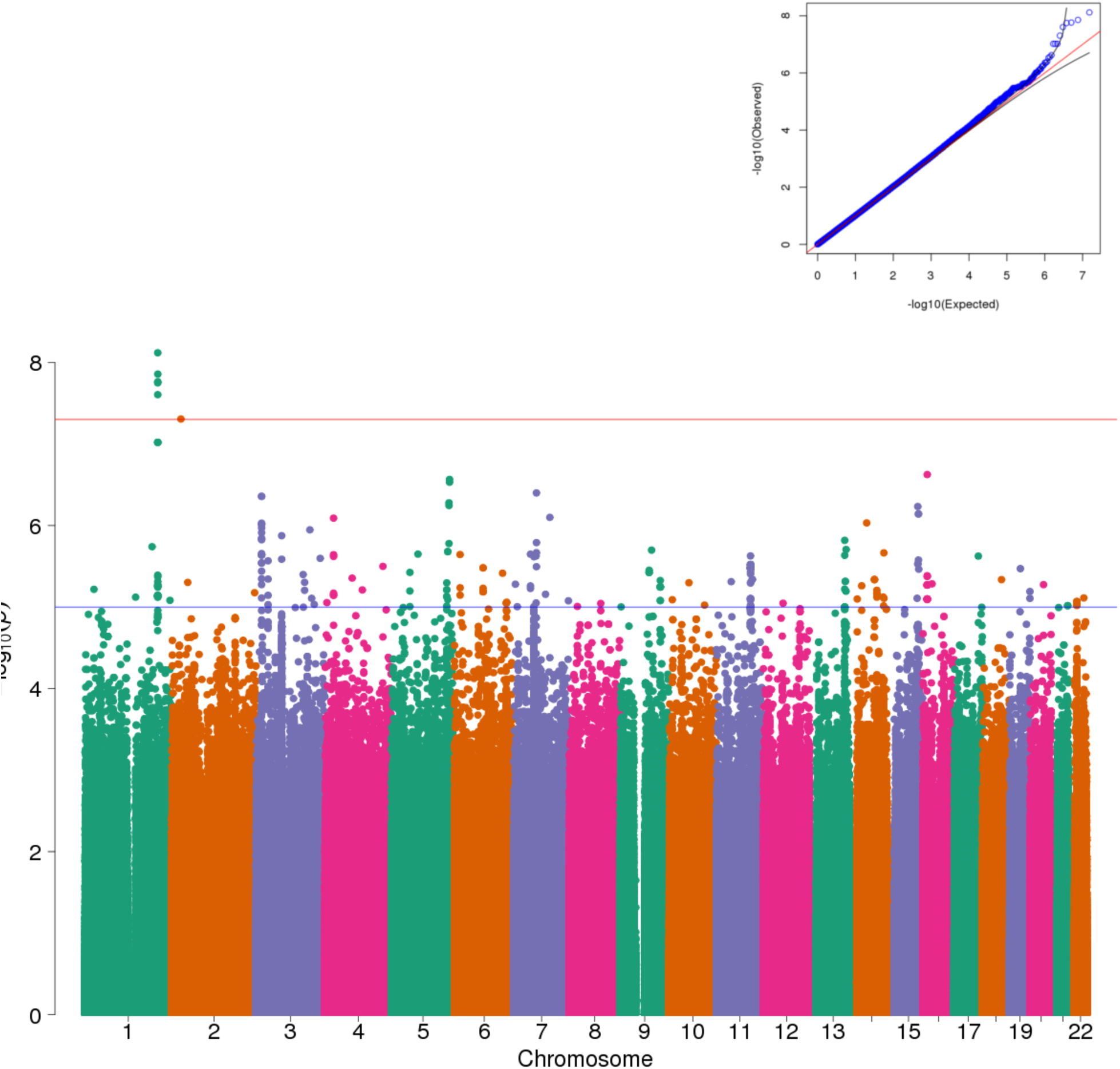
Manhattan plot of the genome-wide association study performed on Δ SBP with Hispanic/Latinos. The top SNP is rs6657443, located on chromosome 1, at locus 209189265 (in hg19 coordinates). The minor allele frequency is 7.63 % in HL (8.83% in the whole cohort), the imputation info score is 0.948. The association p-value is 7.62e-9. Lambda-gc = 1.0027.

**Figure 4A.**
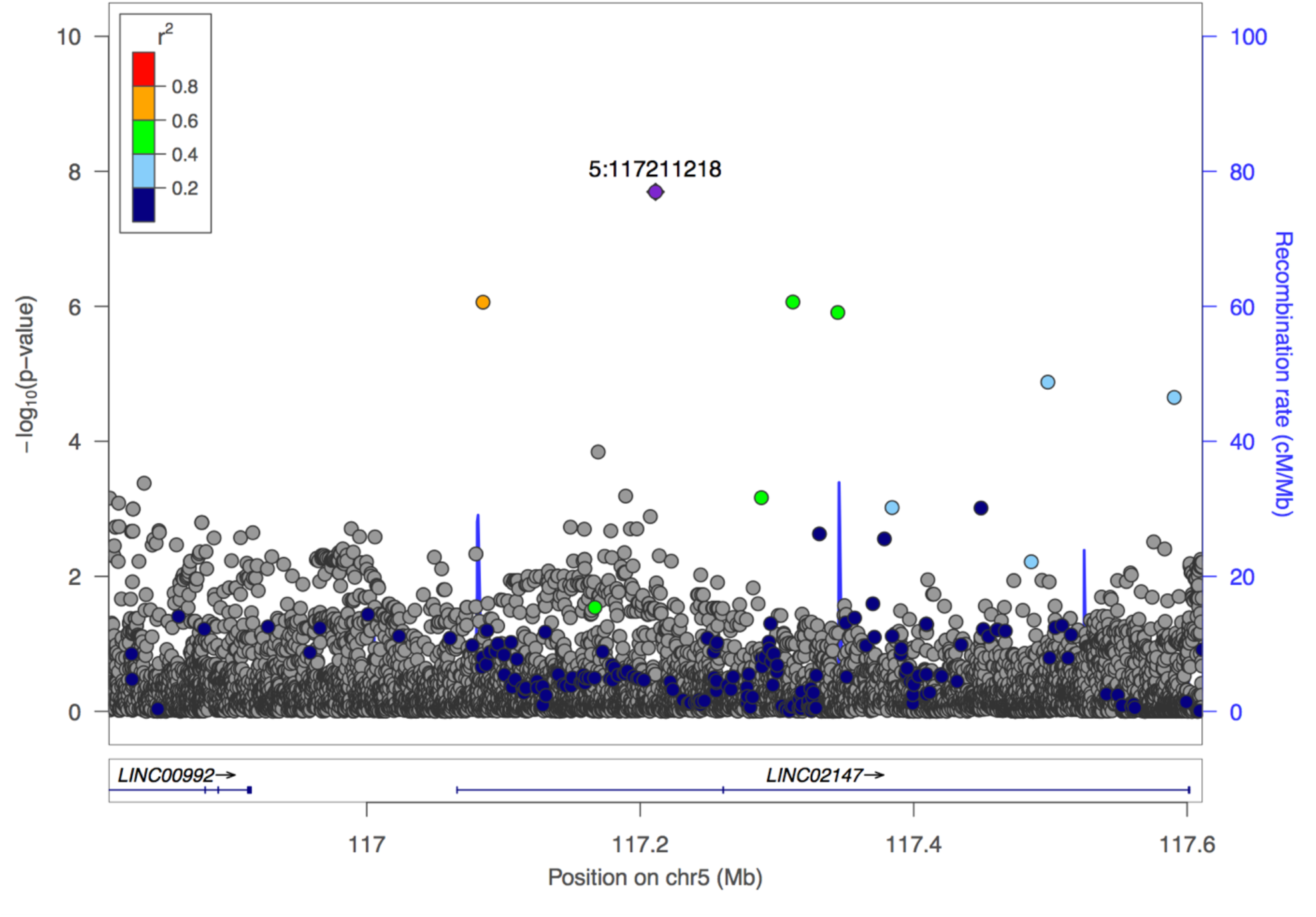
Locus Zoom plot showing patterns of linkage disequilibrium around the top SNP rs145337816 for the genome-wide association study performed on Δ SBP with the whole cohort.

**Figure 4B.**
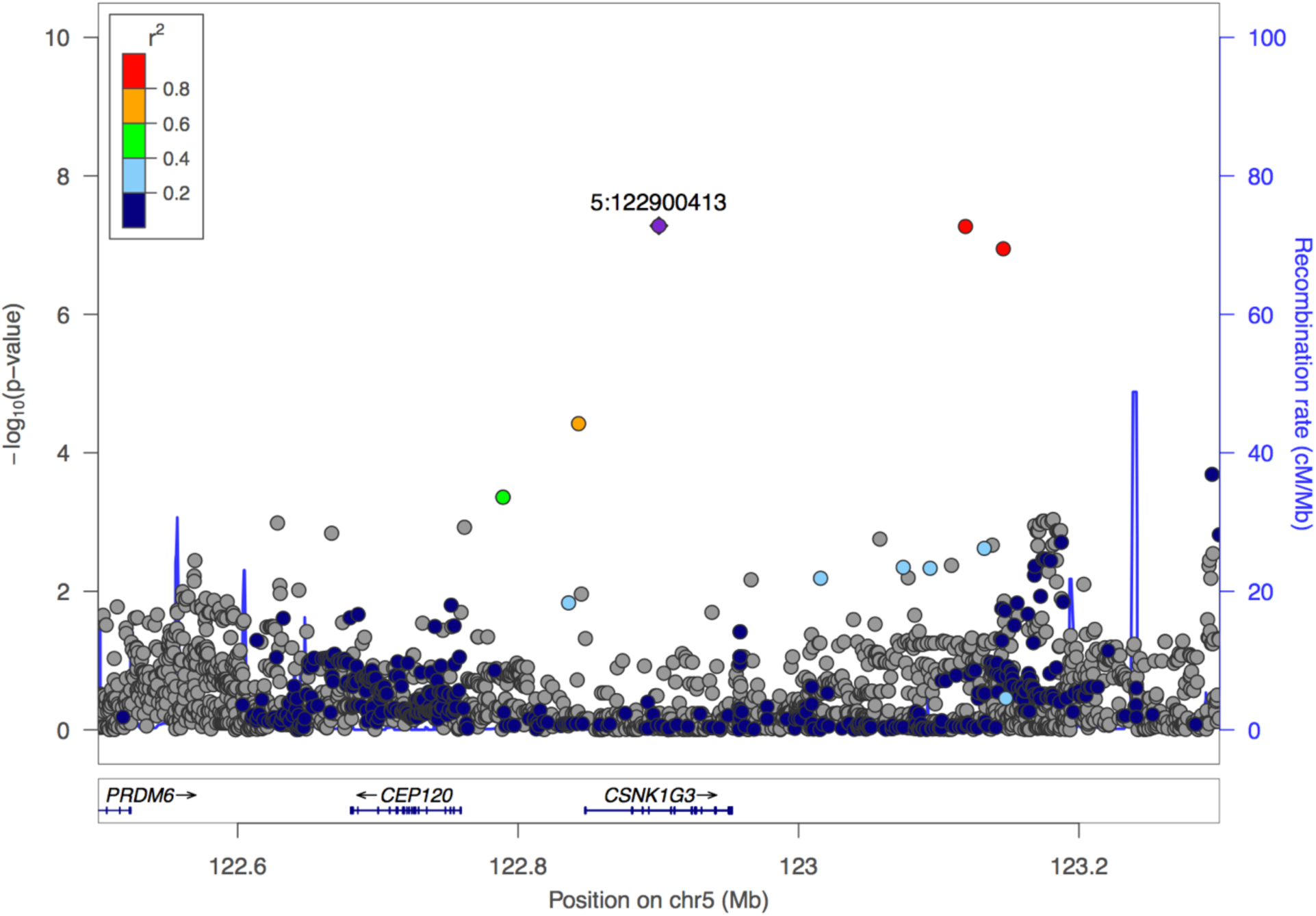
Locus Zoom plot showing patterns of linkage disequilibrium around the top SNP rs188427942 for the genome-wide association study performed on Δ SBP with European Americans.

**Figure 4C.**
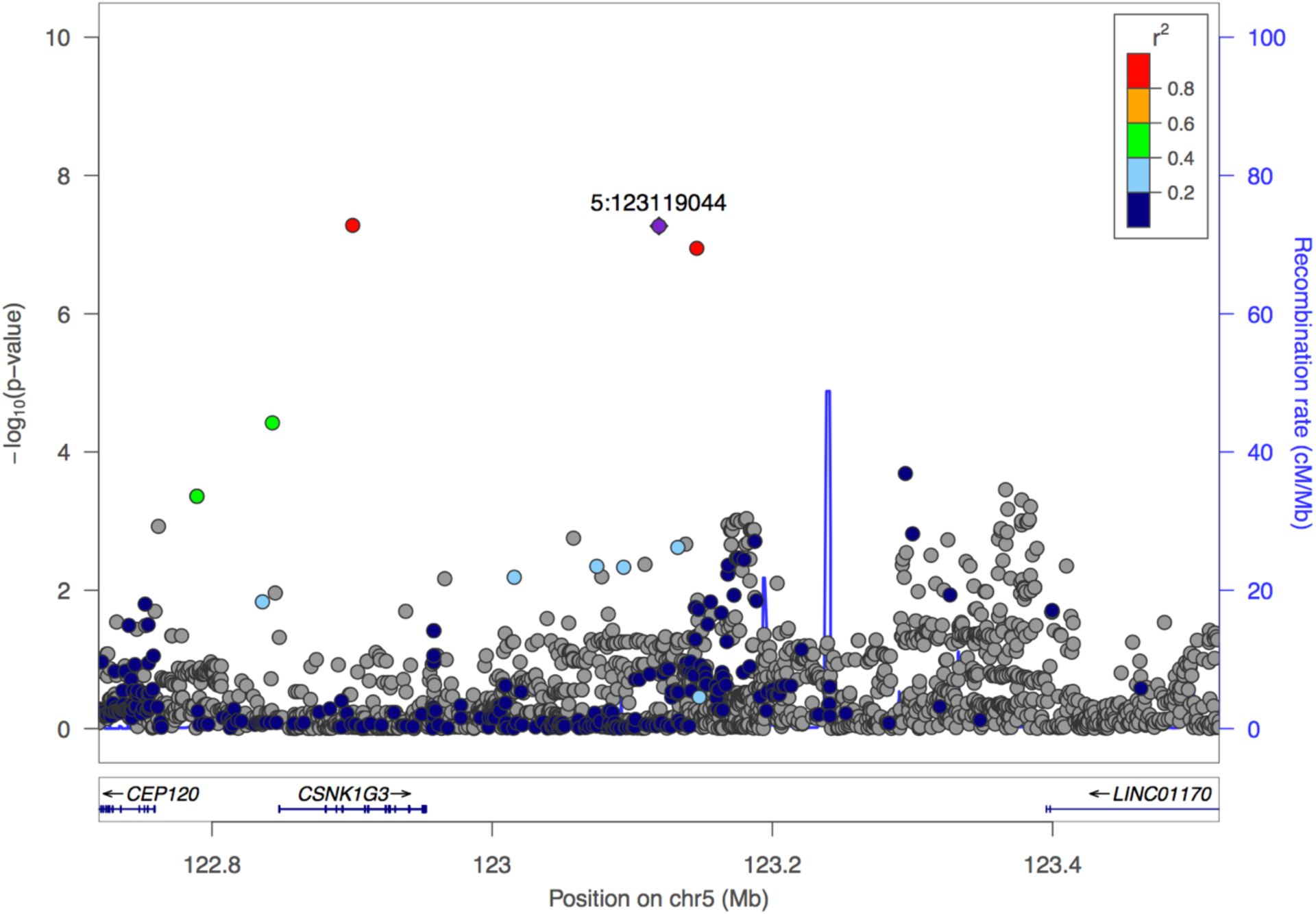
Locus Zoom plot showing patterns of linkage disequilibrium around SNP rs147664194 for the genome-wide association study performed on Δ SBP with European Americans.

**Figure 4D.**
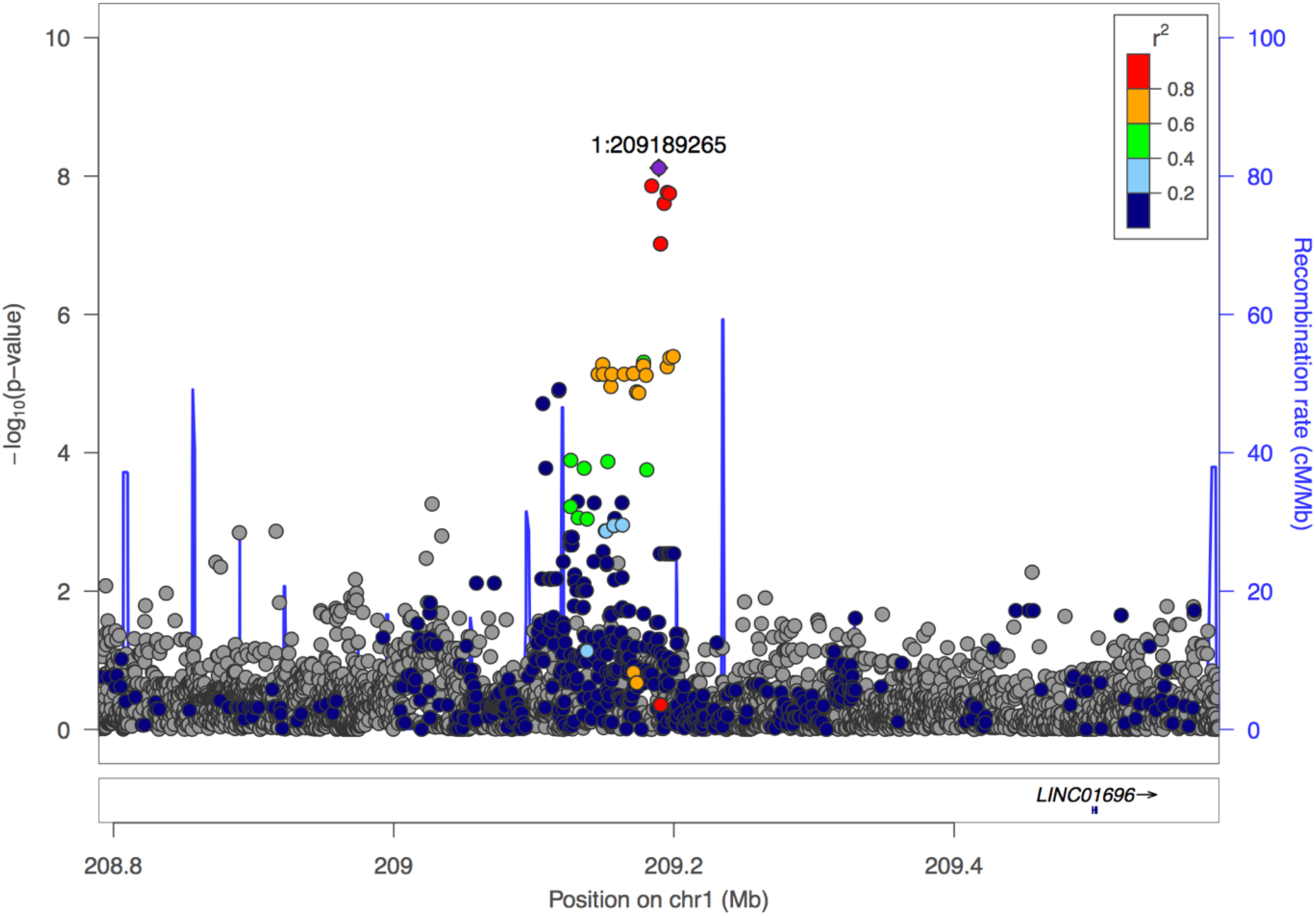
Locus Zoom plot showing patterns of linkage disequilibrium around the top SNPs for the genome-wide association study performed on Δ SBP with Hispanic/Latinos.

**Figure 5.**
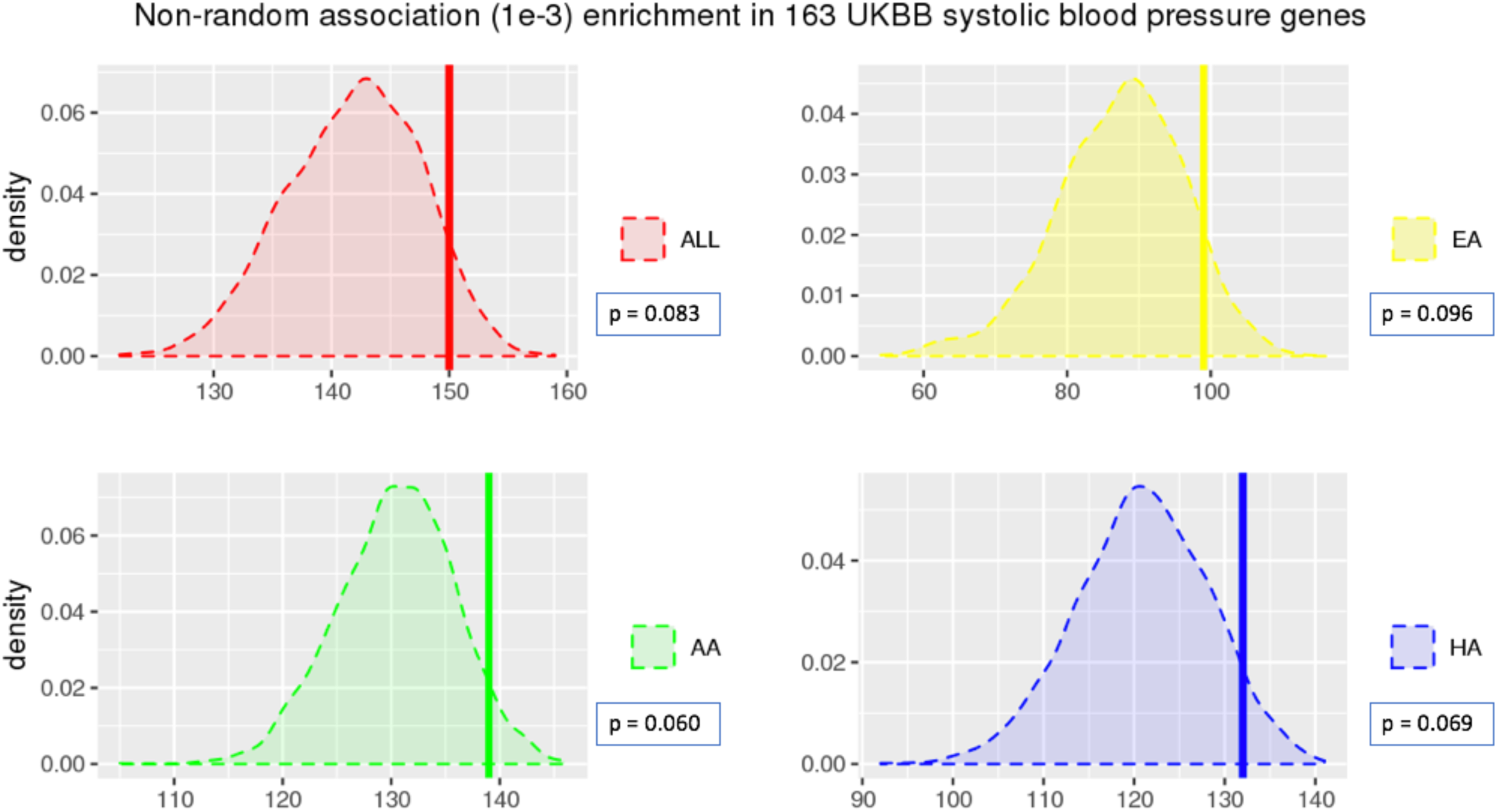
Association enrichment in systolic blood pressure genes. The 4 plots show the distribution of the number of SNP associations with a p-value lower than 1e-3, when randomly selecting 165 genome regions (1000 permutations) for the 4 different population groups (the whole cohort in black, European Americans in green, African Americans in red, Hispanic/Latinos in blue). The vertical bars correspond to the number of genes out of the 165 genes identified in the UKBB cohort to be associated with systolic blood pressure which obtain a p-value lower than 1e-3 when looking for association with Δ SBP in our 4 cohorts. The 165 systolic blood pressure genes are nominally enriched in association with Δ SBP in European Americans (p-value = 0.096), Hispanic/Latinos (p-value = 0.069), African Americans (p-value = 0.060) and in the whole cohort (p-value = 0.083).

## Results

### Validation of phenotype modelling

The initial cohort of 19,685 iGAS patients were matched to 55,104 procedures. After all exclusion steps, our clinical cohort contained a total of 4,317 patients (1,217 AA, 1,387 EA, 1,713 HL). Demographics for the clinical cohort are shown in **Table 1**. Among the patients with valid clinical data, 3,699 were genotyped and passed QC metrics (1,082 AA, 1,112 EA, 1505 HL). (**Figure 1B**). The average values over the entire cohort for the three different BP response measures (Δ SBP, DBP and MAP) were Δ SBP = 17 mmHg (± 25), Δ MAP = 14 mmHg (± 18), Δ DBP = 11 mmHg (± 14). To validate our method of extraction, we compared the observed drug response in our cohort with published estimates from Schwinn et. al. (Schwinn & Reves, 1989). Schwinn performed a small clinical trial on phenylephrine drug response in Europeans and reported a Δ MAP of ∼15 mmHg after 100µg bolus of phenylephrine. Our results were similar with an observed Δ MAP of 15.8 mmHg in European Americans. Likewise, for the entire cohort (all ancestries) we observe an average Δ MAP of 13.9 mmHg. (**Table 2**).

**Table 1.**
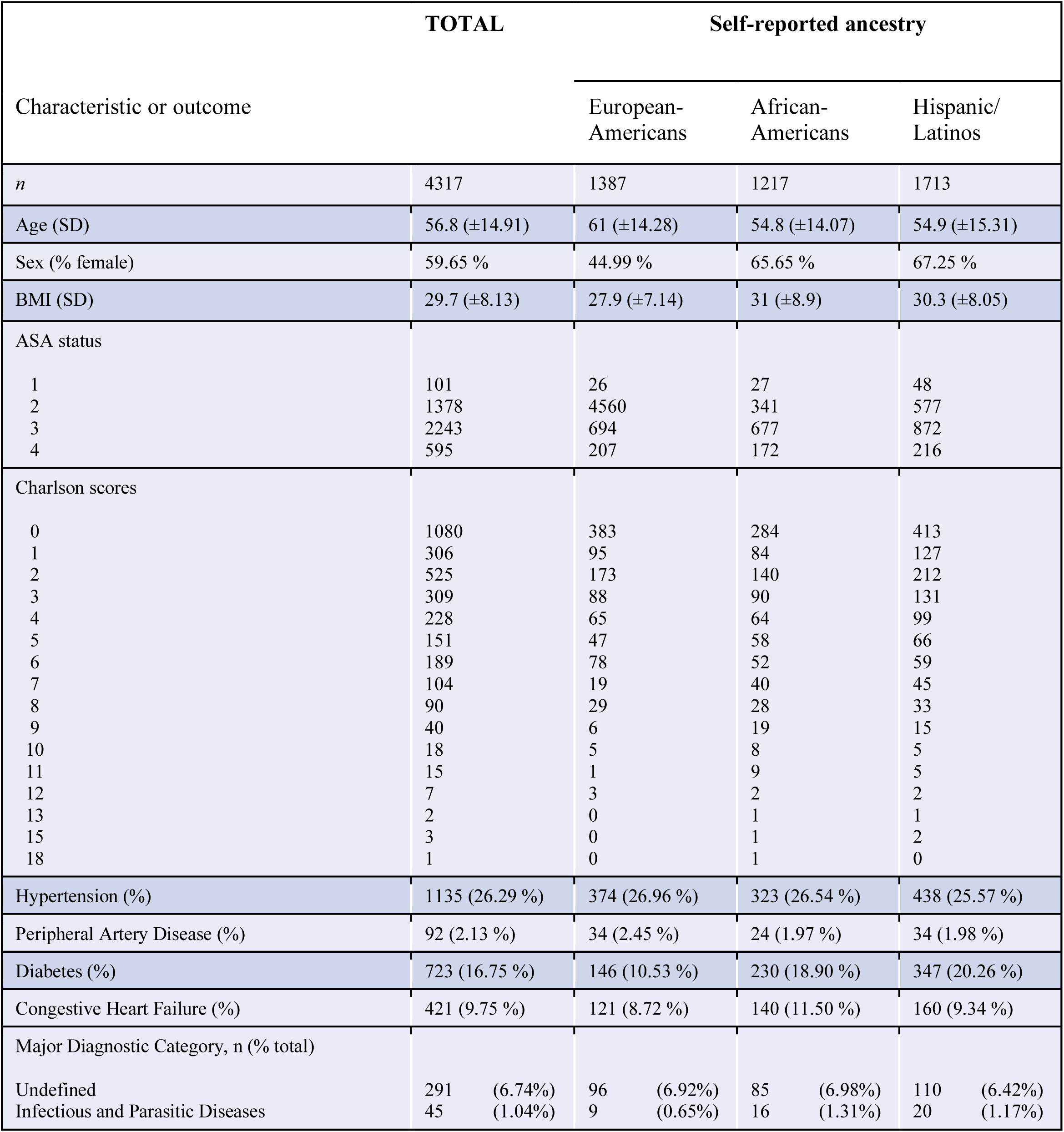
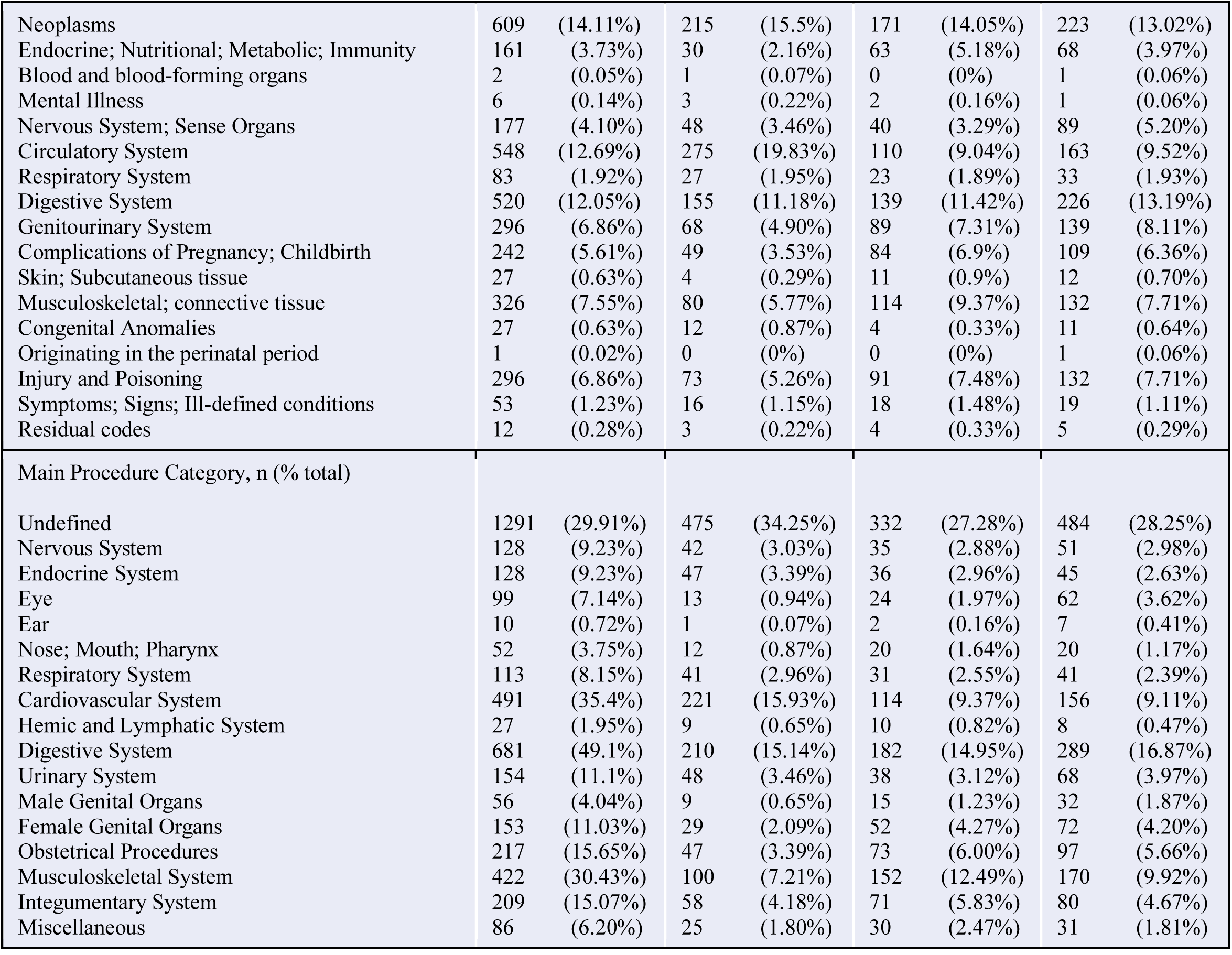
Demographic characteristics of the cohort used in the study.

**Table 2.** Intra-operative characteristics of the cohort used in the study.

| Characteristic or outcome | TOTAL | Self-reported ancestry |  |  |
| --- | --- | --- | --- | --- |
|  |  | European-Americans | African-Americans | Hispanic/Latinos |
| *n* | 4317 | 1387 | 1217 | 1713 |
| Anesthesia type |  |  |  |  |
| General | 2796 | 917 | 779 | 1100 |
| Mac | 928 | 270 | 266 | 392 |
| Regional | 593 | 200 | 172 | 221 |
| Anesthesia time (min) | 187.9 (±147.8) | 205.0 (±143.5) | 179.6 (±135.2) | 179.9 (±158.3) |
| Procedure time (mins) | 114.1 (±96) | 134.9 (±107.4) | 105.5 (±86.8) | 103.9 (±90.0) |
| Number of boluses per case | 4.4 (±4.08) | 4.9 (±5.0) | 4.1 (±3.2) | 4.2 (±3.7) |
| Number of cases per patient | 1.4 (±0.9) | 1.4 (±0.9) | 1.4 (±0.9) | 1.4 (±0.8) |
| Mean bolus amount (micrograms) | 107.2 (±44.5) | 106.05 (±46.4) | 108.8 (±45.9) | 107 (±41.8) |
| Median bolus amount (micrograms) | 100 | 100 | 100 | 100 |
| Cases with Propofol infusion (%) | 1032 (23.90%) | 326 (23.50%) | 302 (24.82%) | 404 (23.58%) |
| Cases with remifentanyl infusion (%) | 390 (9.03%) | 132 (9.52%) | 111 (9.12%) | 147 (8.58%) |
| Cases with phenylephrine infusion (%) | 179 (4.15%) | 82 (5.91%) | 39 (3.2%) | 58 (3.39%) |
| Propofol infusion rate (mcg/kg/min) | 21 (± 43.1) | 20 (±41.5) | 22 (±44.2) | 21 (±43.5) |
| Remifentanyl infusion rate (mcg/kg/min) | 0.01 (±0.13) | 0.01 (±0.07) | 0.01 (±0.05) | 0.01 (±0.19) |
| Phenylephrine infusion rate (mcg/kg/min) | 0.02 (±0.35) | 0.03 (±0.54) | 0.01 (±0.06) | 0.02 (±0.26) |
| Total crystalloids administered (ml) | 1057 (±1031.9) | 1151 (±1082.9) | 1020 (±1017.4) | 1009 (±994.7) |
| Δ Systolic Arterial Pressure, mmHg (SD) | 17.2 (±24.8) | 20.1 (±24.5) | 15.31 (±24.88) | 16.35 (±24.89) |
| Δ Mean Arterial Pressure, mmHg (SD) | 13.9 (±17.7) | 15.8 (±17.4) | 12.8 (±18.0) | 13.2 (±17.7) |
| Δ Diastolic Arterial Pressure, mmHg (SD) | 10.7 (±14.4) | 12.1 (±13.8) | 10.8 (±14.9) | 10.0 (±14.5) |
| Δ Systolic Mann-Whitney vs. EA p-value |  |  | 5.42e-08 | 4.32e-05 |
| Δ MAP Mann-Whitney vs. EA p-value |  |  | 3.52e-06 | 2.35e-04 |
| Δ Diastolic Mann-Whitney vs. EA p-value |  |  | 4.79e-05 | 3.14e-04 |

### Ethnicity impacts BP measures of rapid response to phenylephrine

In a population-stratified analysis, we observed that EA participants demonstrated a significantly heightened rapid response to phenylephrine compared to non-EA patients for all three measures (**Table 2** and **Figure 2)**. The largest difference between populations was observed for Δ SBP (EA Δ SBP = 20 mmHg ± 24; HA Δ SBP = 16 mmHg ± 25; AA Δ SBP = 15 mmHg ± 25), thus this phenotype was used for all downstream analysis. After adjusting for age, sex, BMI, ASA status, phenylephrine bolus amount, depth of anesthesia, total volume of fluid administered, and accounting for genetic ancestry, significant differences remained for all three measures between EA participants and HA (Δ SBP, p < 0.032; Δ MAP, p < 0.021; Δ DBP, p < 0.008), and between EA and AA participants (Δ SBP, p < 5.13e-5; Δ MAP, p < 2.1e-4; Δ DBP, p < 3.3e-4).

### Genetic Discovery

We performed genome-wide association studies for systolic blood pressure response to phenylephrine (Δ SBP), as well as Δ MAP and Δ DBP, across > 47 million SNPs either imputed from 1,000 genomes project or genotyped directly in 3699 individuals (1082 AA, 14505 HA, 1112 EA) altogether, as well as for each of the 3 populations separately. Across the three phenotypes (Δ SBP, Δ MAP, Δ DBP) and the four population groupings (the whole cohort, African Americans, European Americans, Hispanic/Latinos) we observed genome-wide significant associations (p-value < 5e-8) in 5 different genome regions. Additionally, we observed 2 genome regions with suggestive associations at p-values < 6e-8 and < 7e-8 in EAs. Among these 7 regions, 5 of them overlap variants previously associated with routine blood pressure measurements (**Table 3, Figures 3-4, Supplementary Figures 7-9**). Remarkably, the analysis performed on the HA population with the Δ SBP phenotype identified five adjacent genome-wide significant SNPs which are common in all three studied populations (all MAF > 7%), yet these variants are only associated with a significantly decreased systolic response in the HA population, thereby asserting the importance of ancestry as a factor influencing common alleles’ effect size. Notably, these five SNPs are located at 1.27 Mb upstream of rs2761436, a variant previously associated with SBP (Hoffmann et al., 2017) (**Table 3, Figure 3, Figure 4**).

**Table 3.** Results of the genome-wide association studies performed with the three different phenotypes (Δ SBP, Δ MAP, Δ DBP) and the four population groupings (the whole cohort, African Americans, European Americans, Hispanic/Latinos). The average carrier phenotypes are shown only for SNPs with 10 carriers or more. SNPs previously associated with BP phenotypes in other studies are reported if they’re located within a 1 Mb genomic region overlapping the corresponding hit SNP, with the exception of rs2761436 (1.27 Mb). ^1^: PP: pulse pressure. AT: arterial tonometry. (Andreassen et al., 2014; Ehret et al., 2016; Evangelou et al., 2018; Giri et al., 2019; Kichaev et al., 2019; Levy et al., 2007; Liu et al., 2016; Nielsen et al., 2018; Roselli et al., 2018; Salvi et al., 2017; Taylor et al., 2016)

| Phenotype | Population | Chr | BP (hg19) | Beta | SE | P-value | Associated gene(s) | rsid | Average Carrier Phenotype (mmHg) |  |  | MAF |  |  |  | Previous associations with BP phenotype<sup>1</sup> |
| --- | --- | --- | --- | --- | --- | --- | --- | --- | --- | --- | --- | --- | --- | --- | --- | --- |
|  |  |  |  |  |  |  |  |  | EA<br>n = 1112 | AA<br>n = 1082 | HA<br>n = 1505 | EA<br>n = 1112 | AA<br>n = 1082 | HA<br>n = 1505 | ALL<br>n = 3699 |  |
| SBP | ALL | 5 | 117,211,218 | 0.92 | 0.16 | 2.02E-08 | LINC02147 | rs145337816 | 37.3 +- 29.1 | - | 53.1 +- 36.2 | 0.56% | 0.00% | 0.33% | 0.46% |  |
|  | EA | 5 | 122,900,413 | -1.10 | 0.20 | 5.26E-08 | CSNK1G3 | rs188427942 | -6.8 +- 30.4 | - | 6.9 +- 42.3 | 1.16% | 0.28% | 0.34% | 0.57% | rs1579036 (DBP),<br>rs4530754 (PP),<br>rs11750782 (SBP),<br>rs373496042 (SBP),<br>rs6891344 (DBP, SBP) |
|  |  | 5 | 123,119,044 | -1.17 | 0.22 | 5.38E-08 |  | rs147664194 | -9.1 +- 31.6 | - | - | 1.01% | 0.23% | 0.27% | 0.48% |  |
|  | HA | 1 | 209,189,265 | -0.39 | 0.07 | 7.62E-09 | - | rs6657443 | 21.0 +- 28.0 | 14.0 +- 27.1 | 7.3 +- 24.0 | 7.87% | 11.59% | 7.63% | 8.83% | rs2761436 (SBP) |
|  |  | 1 | 209,184,198 | -0.36 | 0.06 | 1.39E-08 |  | rs150793135 | 20.4 +- 27.6 | 14.2 +- 26.3 | 8.3 +- 23.7 | 7.80% | 16.62% | 9.03% | 10.77% |  |
|  |  | 1 | 209,195,248 | -0.39 | 0.07 | 1.72E-08 |  | rs6670957 | 21.0 +- 28.0 | 13.6 +- 26 | 7.2 +- 23.6 | 7.87% | 10.29% | 7.17% | 8.28% |  |
|  |  | 1 | 209,196,851 | -0.40 | 0.07 | 1.78E-08 |  | rs34954876 | 21.1 +- 28.2 | 13.6 +- 26 | 7.2 +- 23.7 | 7.76% | 10.34% | 7.14% | 8.24% |  |
|  |  | 1 | 209,193,053 | -0.39 | 0.07 | 2.49E-08 |  | rs139349180 | 21.0 +- 28.0 | 14.0 +- 26.5 | 7.4 +- 23.7 | 7.87% | 10.40% | 7.21% | 8.32% |  |
| MAP | ALL | 13 | 30,950,119 | 0.14 | 0.02 | 1.97E-08 | LINC01058,<br>LINC00426,<br>KATNAL1, HMGB1 | rs145222507 | 16.6 +- 18.0 | 13.0 +- 17.4 | 15.3 +- 17.8 | 33.96% | 39.10% | 32.34% | 34.82% |  |
|  |  | 13 | 30,944,206 | 0.13 | 0.02 | 2.81E-08 |  | rs1275187 | 16.0 +- 18.3 | 13.1 +- 17.5 | 15.1 +- 17.9 | 34.06% | 39.72% | 32.42% | 35.05% |  |
|  |  | 13 | 30,948,845 | 0.13 | 0.02 | 2.85E-08 |  | rs1275192 | 16.0 +- 18.2 | 13.1 +- 17.3 | 15.3 +- 17.8 | 34.52% | 39.81% | 33.27% | 35.50% |  |
|  |  | 13 | 30,946,291 | 0.13 | 0.02 | 3.53E-08 |  | rs1275189 | 16.0 +- 18.3 | 13.1 +- 17.6 | 15.1 +- 17.8 | 33.86% | 39.64% | 32.32% | 34.92% |  |
|  |  | 13 | 30,946,686 | 0.13 | 0.02 | 3.81E-08 |  | rs1275191 | 16.0 +- 18.3 | 13.1 +- 17.6 | 15.1 +- 17.8 | 33.97% | 39.64% | 32.34% | 34.97% |  |
|  |  | 13 | 30,945,458 | 0.13 | 0.02 | 4.17E-08 |  | rs21498505458 | 16.0 +- 18.3 | 13.1 +- 17.5 | 15.1 +- 17.8 | 34.06% | 39.72% | 32.39% | 34.94% |  |
|  | EA | 1 | 41,519,758 | 0.49 | 0.09 | 6.38E-08 | SCMH1, SLFNL1-AS1, SLFNL1,<br>CTPS1 | rs111908123 | 23.8 +- 19.2 | 13.0 +- 20.8 | 14.5 +- 18.8 | 5.90% | 1.22% | 4.25% | 3.82% | rs11210029 (SBP) |
| DBP | AA | 8 | 16,559,461 | -1.27 | 0.22 | 1.13E-08 | - | rs146535276 | - | -8.1 +- 16.8 | 11.4 +- 19.4 | 0.05% | 0.99% | 0.33% | 0.44% | rs11784910 (SBP,<br>DBP), rs3793427 (AT),<br>rs75902664 (DBP) |
|  | AA | 16 | 59,684,462 | 1.24 | 0.22 | 1.93E-08 | APOOP5 | rs143947120 | - | 27.5 +- 18.2 | - | 0.00% | 0.99% | 0.27% | 0.40% | rs182346790 (PP),<br>rs35300112 (SBP),<br>rs1862746 (SBP) |

We identified two “non-responder” groups: patients who either had no increase in blood pressure in response to the bolus, or whose blood pressure paradoxically declined. The first group of non-responders was identified through the analysis performed on the EA population, which identified two rare SNPs, rs188427942 (MAF EA: 1.16%) and rs147664194 (MAF EA: 1.01%) located in or near gene CSNK1G3 (Casein Kinase 1 Gamma 3), in regions previously associated with SBP, DBP, and PP (pulse pressure) (Andreassen et al., 2014; Ehret et al., 2016; Giri et al., 2019; Liu et al., 2016). CSNK1G3, which encodes a member of a family of serine/threonine protein kinases that phosphorylate caseins and other acidic proteins, is mostly expressed in fibroblasts (GTEx Consortium, 2013) (**Supplementary Figure 6**). On average, the 22 carriers of both SNPs have a Δ SBP of -9 mmHg (± 32). In contrast the rest of the EA samples had an average Δ SBP of 20 mm Hg (± 24). The second group of non-responders was identified through the analysis performed on the AA population with the Δ DBP phenotype. Intergenic SNP rs146535276 (MAF AA: 0.99%) is located in a 1 Mb genomic region on chromosome 8 previously associated with SBP, DBP, and AT (arterial tonometry) (Evangelou et al., 2018; Levy et al., 2007; Salvi et al., 2017). On average, the 21 AA carriers of rs146535276 have a Δ SBP of -2 mmHg (± 26) mmHg, while the rest of the AA cohort have a Δ SBP of 15 mmHg (± 25) (**Table 3, Figure 3, Figure 4, Supplementary Figure 9**). Remarkably, we also observed that carriers of rs145337816, a rare SNP only present in European Americans (MAF: 0.56%) and Hispanic/Latinos (MAF: 0.33%), constitute a small group of potential “super-responders”, i.e. patients with a significantly high response (Δ SBP_EA_= 37 mmHg (± 29) and Δ SBP_HA_= 53 mmHg (± 36)), with the caveat that results obtained on such a small number of individuals might not be reproducible.

### Systolic Blood Pressure Genes Association

We observed non-random enrichment in association (significance threshold set at p < 0.001) with systolic drug response in 165 loci previously reported to be associated with systolic blood pressure in the UK BioBank cohort. These 165 loci are centered around single nucleotide polymorphisms (SNPs) reported as being associated (p < 5e-8) with systolic blood pressure in a genome-wide association study performed on 361,402 individuals of mostly European ancestry (Sudlow et al., 2015). To explore whether the genetic loci underlying SBP as measured in routine clinical visits may impact the Δ SBP response to phenylephrine administration during surgery, we investigated whether these loci were enriched in the GWAS signals. First, we counted the number of times we observed a p-value of association less than 10-3 in all 165 regions. Next, we counted the number of times we observed a p-value of association less than 10-3 in 1000 random selections of 165 regions of similar genomic size, in order to generate a distribution of random signal across the genome (**Figure 4**). Out of the 165 independent genome regions associated with SBP in the UK BioBank cohort, 150, 139, 99 and 132 regions were associated with Δ SBP in our complete cohort, and in AA, EA and HA individuals respectively. Alternatively, when randomly selecting 165 regions of the genome, 141.9 (± 5.69), 130.4 (± 5.41), 87.4 (± 8.95), 120.9 (± 7.34) regions are associated with Δ SBP in our complete cohort, and in AA, EA, HA individuals respectively (1000 random selections were performed). Comparing these random distributions with the ratio of systolic blood pressure regions associated with Δ SBP in the 4 cohorts allows us to establish p-values of 0.096, 0.069 and 0.06 for EA, HA, and AA participants respectively and of 0.093 for the full cohort. These results indicate a shared genetic architecture underlying both phenylephrine response and systolic blood pressure.

## Discussion

The past decade has seen tremendous growth in the use of clinical data from the Electronic Health Record (EHR) for genomic discovery (Bowton et al., 2014; Kohane, 2011; McCarty et al., 2011). Much, if not all, of this research has focused on longitudinal data; for example, chronic disease conditions recorded in the EHR, or blood pressure taken during routine annual outpatient office visits (Dumitrescu et al., 2017; Hoffmann et al., 2017). These data, while valuable for the observation of phenotypes, are of low temporal resolution. Thus, most phenotypes observed in this manner are best suited for genomic discovery in the context of chronic (i.e., long term) medical conditions. In contrast, as pharmacogenomics has traditionally focused on drugs administered in a controlled environment, there has been limited research to date on the influence that genetics may have on the physiologic response to more acute clinical conditions, such as those experienced during surgery or during a hospital admission.

In the present study, we successfully used extant clinical data recorded during surgery for the secondary purpose of pharmacogenomics research. We underscored the importance of defining a rigorous phenotype to represent the dose-response to phenylephrine during surgery. Reassuringly, the results of our derived phenotype closely matched the BP change observed by Schwinn et. al. 30 years ago (Schwinn & Reves, 1989). This remarkable consistency validates both Schwinn’s original results obtained in a small cohort, as well as our contemporary results derived from routine clinical care. Our phenotypic modelling process allowed us to perform population-level comparisons, notably highlighting statistically significant differences across self-reported ancestries. We found that self-identified Europeans had a significantly heightened response to phenylephrine administered intra-operatively compared to other populations. We tested several hypotheses to explain this difference, including, age, gender, body mass index, level of anesthesia, concurrent medications, and co-morbidities **(Table 1)**. None of the aforementioned variables explained the difference in phenylephrine response across populations. Differences in drug response between populations provide strong evidence that genetic ancestry and/or unmeasured environmental factors could account for the trait variability across populations, as is the case with other drug response or pharmacogenomics phenotypes (Ramos et al., 2014; Wright, Carleton, Hayden, & Ross, 2016).

Using this derived trait in a GWAS allowed us to discover novel loci in the genome that could explain variability in drug response to phenylephrine. Notably, five out of the seven independent genomic regions harboring SNPs associated with the three derived phenotypes were previously associated with related blood pressure phenotypes. Moreover, the enrichment in association with systolic blood pressure genes is indicative of a likely shared genetic architecture between systolic drug response and systolic blood pressure, a commonality previously unknown. Our experimental design allowed us to identify common variants present in all three populations and having significantly different effect sizes on the different populations. Additionally, we identified rare SNPs, in the analyses performed separately on the European American cohort and the African American cohort, whose carriers are phenylephrine non-responders. Being able to predict non-responders based on a small number of SNPs has clinical implications, especially in an era of personalized medicine, where the number of patients being genotyped or sequenced grows exponentially.

These findings confirm an effect that has been anecdotally observed for many years and may have important potential clinical implications. Our study adds to the existing body of knowledge by significantly increasing the number of sampled individuals and assessing Δ MAP, Δ SBP and Δ DBP in patients of diverse ancestries, as the extent to which African Americans and Hispanic/Latinos respond to intravenous phenylephrine administration was previously unknown. While it had previously been observed in small cohorts of patients undergoing cardiac surgery with CPB that carriers of the G894T polymorphism of NOS3 as well as carriers of the insertion/deletion polymorphism of ACE have a significantly heightened vascular responsiveness to phenylephrine (Henrion et al., 1999; Lasocki et al., 2002; Philip et al., 1999), we did not observe this effect in our cohort.

### Limitations

Our approach entails some caveats and should be considered as exploratory science rather than hypothesis-based. Even at the scale of hospital-wide biobanks, the application of stringent sample filtering and removal steps, which are paramount when studying a phenotype measured in the “real-world”, dynamic environment of an operating room, prevents us from reaching sample sizes large enough to replicate this method on less commonly used drugs or to extrapolate a polygenic risk score. Using only a single response does not allow us to build a dose-response curve for each patient and it is possible that if we had chosen a different bolus to analyze, we would have seen somewhat different results. Furthermore, while we did include depth of anesthesia at the time of the phenylephrine bolus as a covariate, the BP response observed was likely also dependent on the degree of surgical stimulation (or lack thereof) and volume status at the time of the bolus. These variables were not captured and remain as unmeasured residual confounders. Thanks to the diversity of our biobank, our multi-ethnic study was balanced enough to allow us to observe significant differences in drug response across self-reported population groups. However, given the expanse of our sample set and the inherent statistical limitations of the methods used, genomic signals recorded should be reckoned as potential association indications requiring further downstream analysis rather than definitive assessments of the functional role of genes mentioned. Cardiac output, which is not routinely measured outside of cardiac surgery operating rooms, is a significant residual confounder of the blood pressure response to any pressor. The incidence of congestive heart failure (CHF), which is usually a low cardiac output state, was significantly higher among African Americans in our cohort. It is possible that this is partially responsible for the blunted SBP response we observed in this group, however a multivariate analysis showed no significant association between CHF and SBP in our cohort. We were able to identify groups of non-responders carrying rare variants, however given the small number of patients carrying these variants, several potential confounders might be at play, and these results should be considered preliminary until confirmed by further prospective studies.

As biobanks and genomic databases are constantly increasing in size, limitations in sample size will gradually disappear. Moreover, federated datasets and data-sharing across multiple institutions are paving the way towards sample sizes orders of magnitude larger than currently available ones, leading to significant increases in statistical power and precision (Abul-Husn & Kenny, 2019; Stark, Dolman, Manolio, & Ozenberger, 2019). Approaches like the one used in this paper, leveraging the union of genomics and extensive clinical data capture, as well as statistical tools borrowed from the field of artificial intelligence to customize treatment, will lead to tailored dosage and anesthetic plans, thereby constituting a true paradigm change in the practice of surgery and making it the next field where patients can expect to benefit from precision medicine.

## Author contributions

SW, JMJ conceived and designed the experiments; performed the analyses and contributed to the writing of the manuscript. TJ, MCY, GMB, AOO, SBE, EPB, OG contributed to the writing of the manuscript. MAL, EEK conceived and designed the experiments; contributed to the writing of the manuscript.

