## Supplementary Material for "Rapid response to the Alpha-1 Adrenergic Agent Phenylephrine in the Perioperative Period is Impacted by Genomics and Ancestry"

**Supplements**

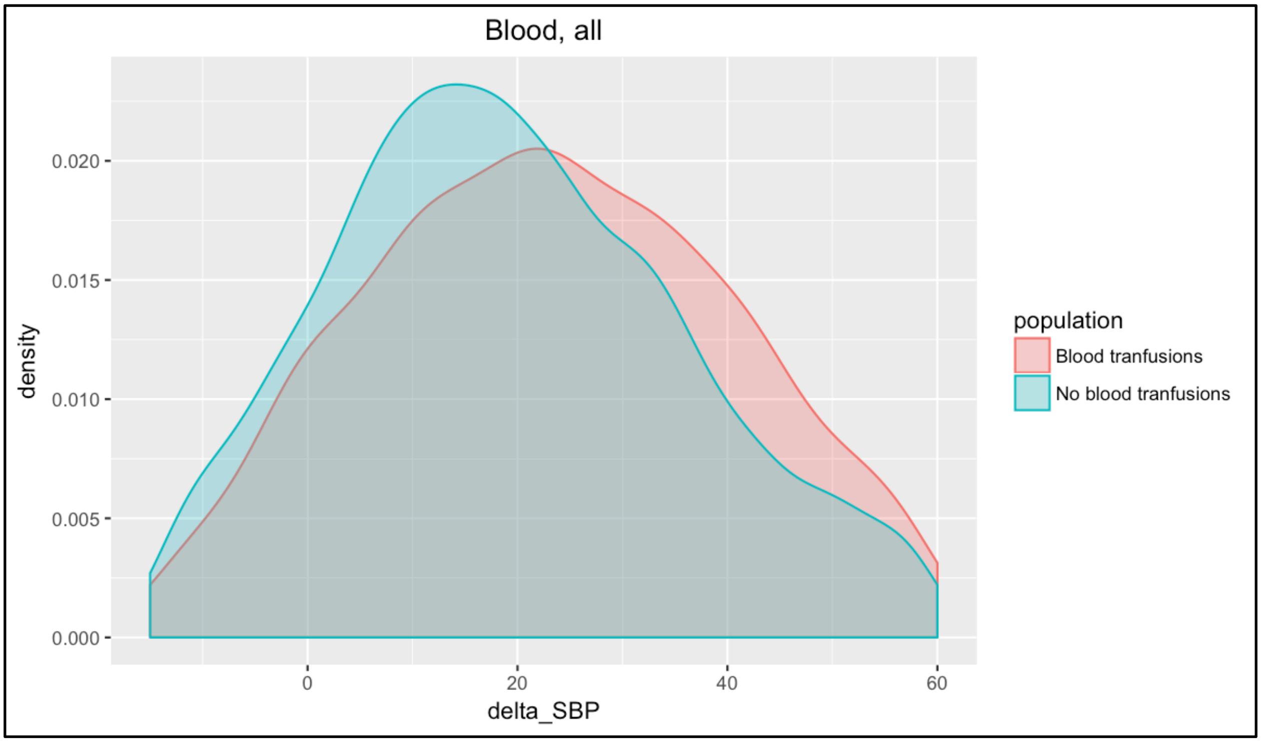


**Supplementary Figure 1.** Phenotype difference between patients having received blood transfusions during surgery and patients without transfusions.

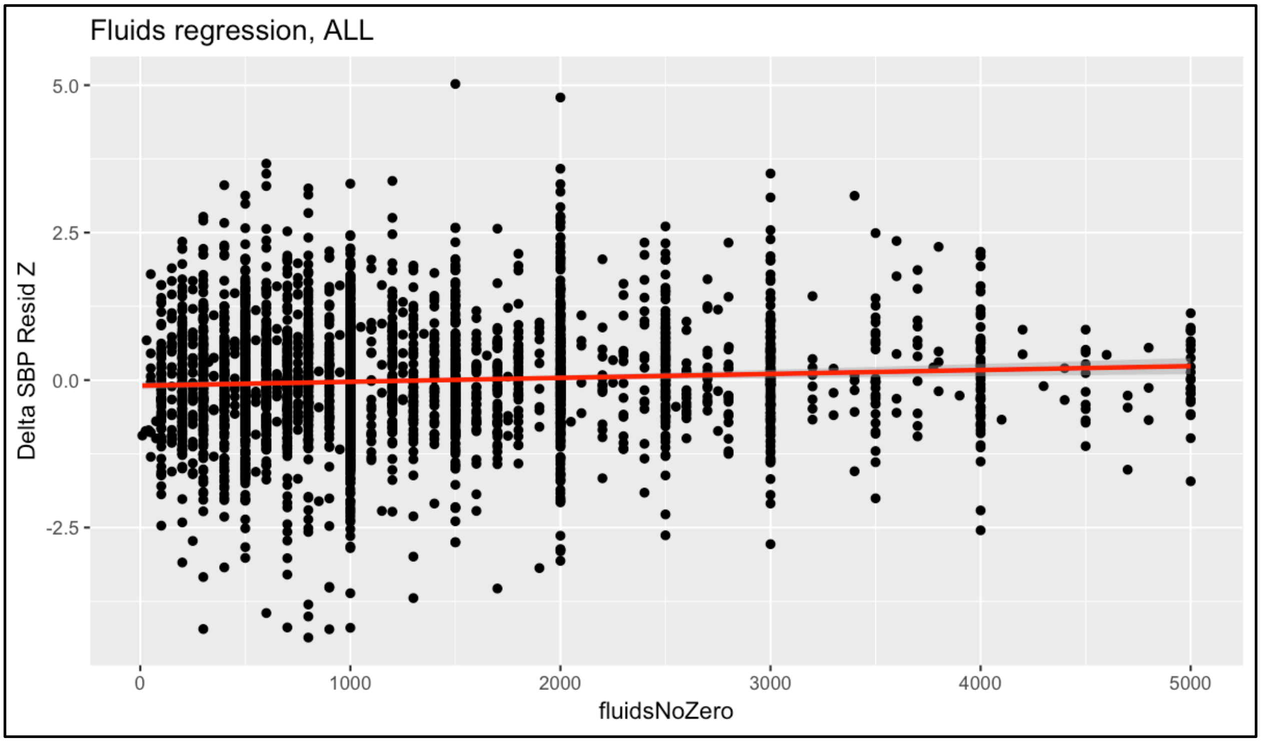


**Supplementary Figure 2.** Association between total amount of crystalloid administered and the phenotype.

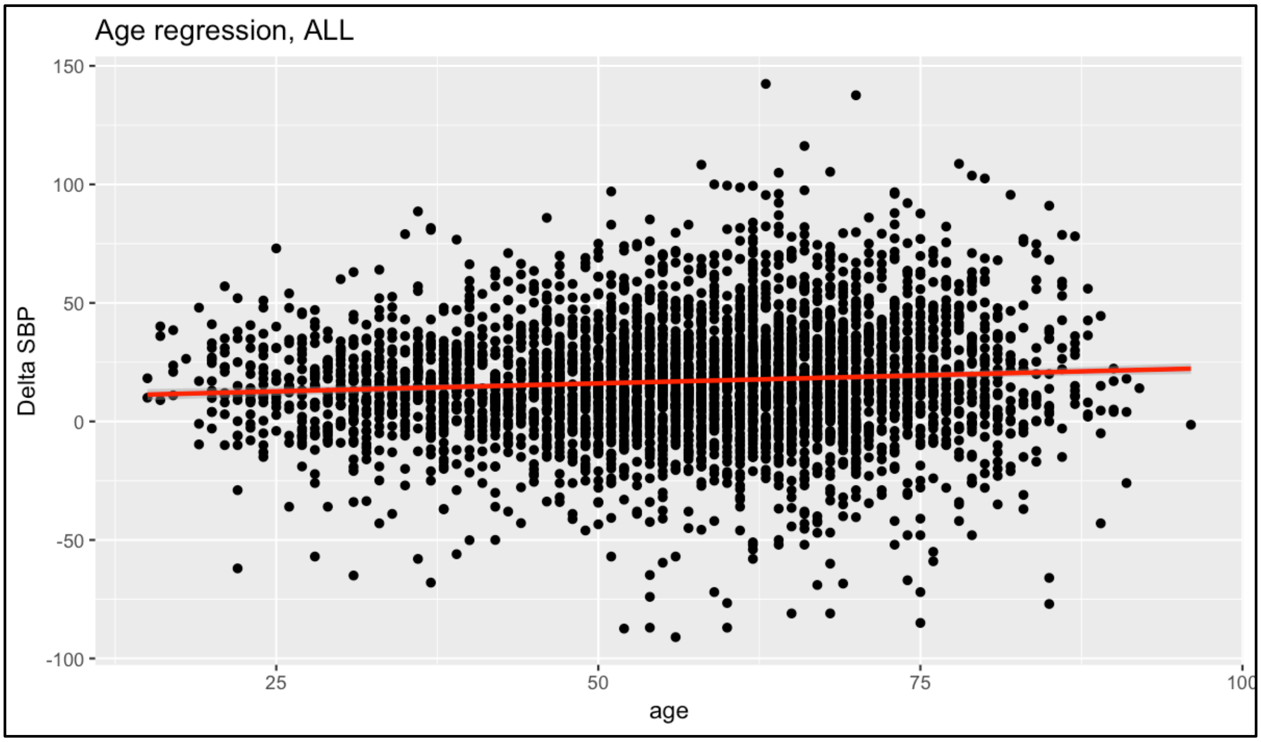


**Supplementary Figure 3.** Association between age and the phenotype.

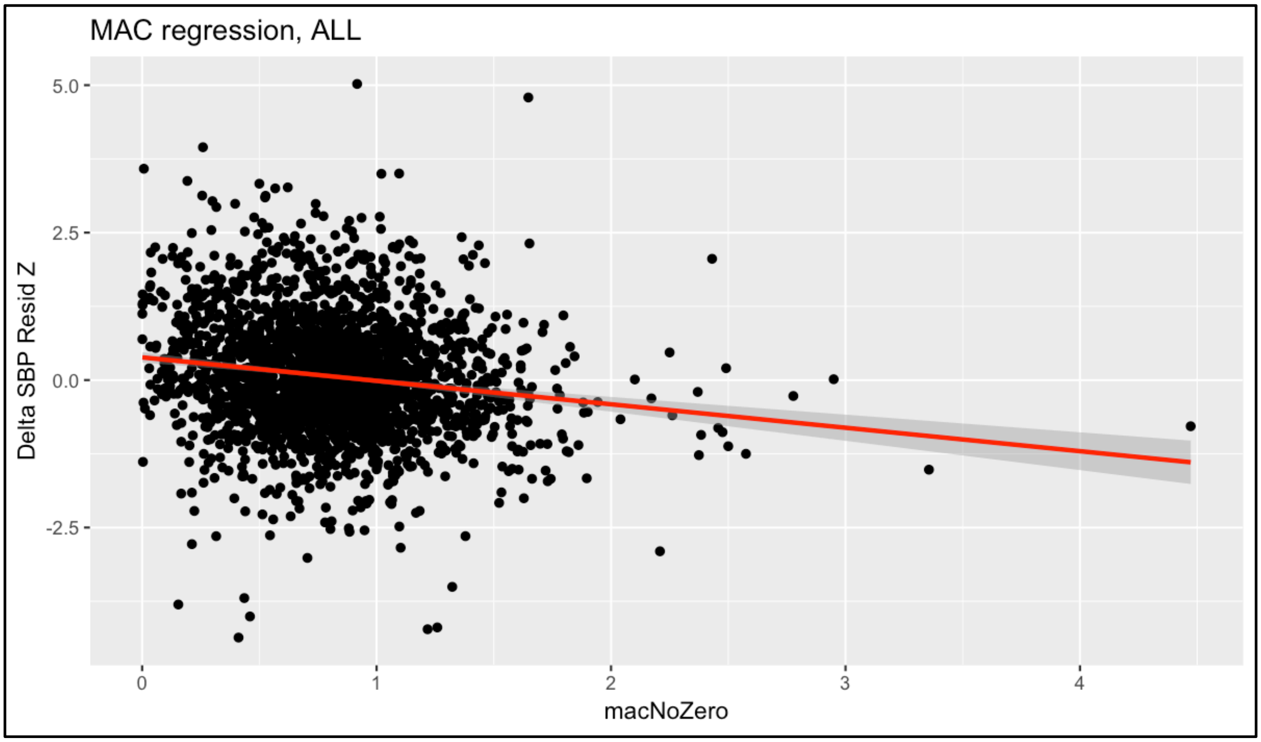


**Supplementary Figure 4.** Association between Minimal Alveolar Concentration and the phenotype.

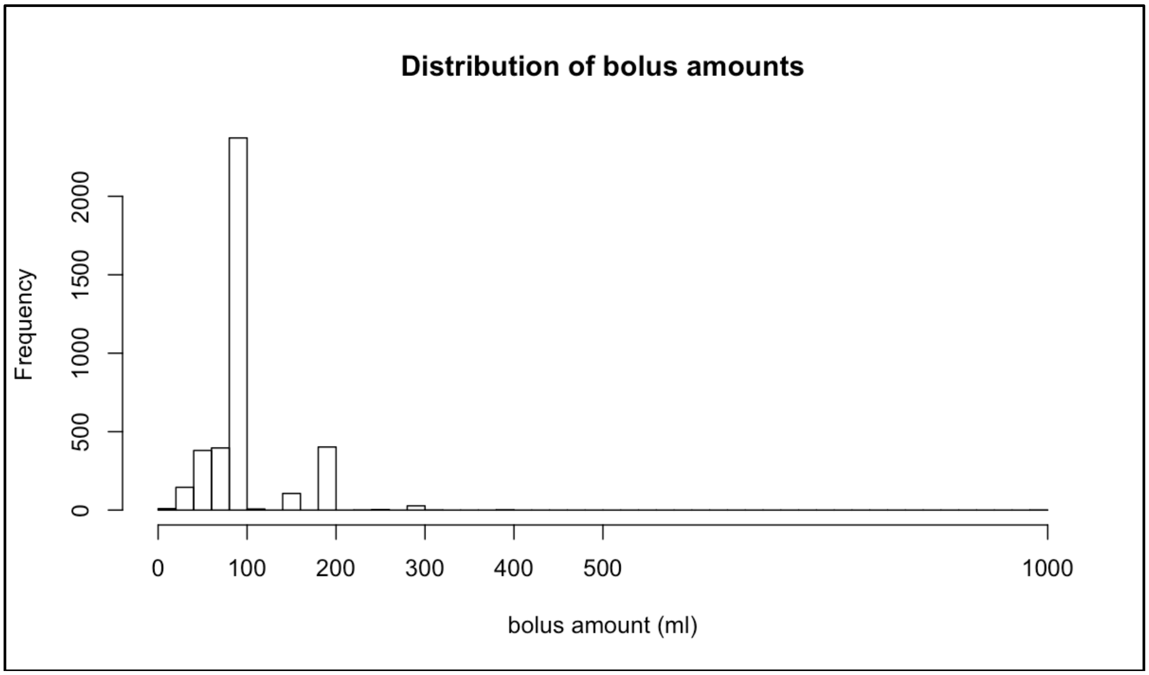


**Supplementary Figure 5.** Distribution of bolus amounts across all patients and all procedures.

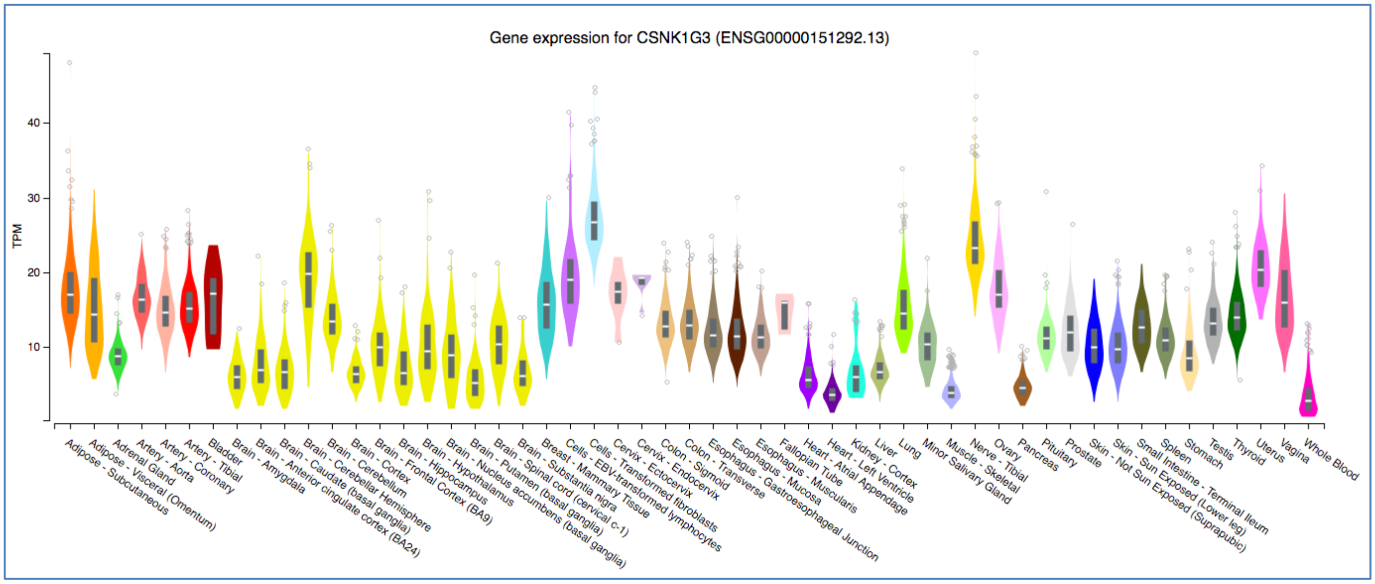


**Supplementary Figure 6.** Tissue expression of CSNK1G3.

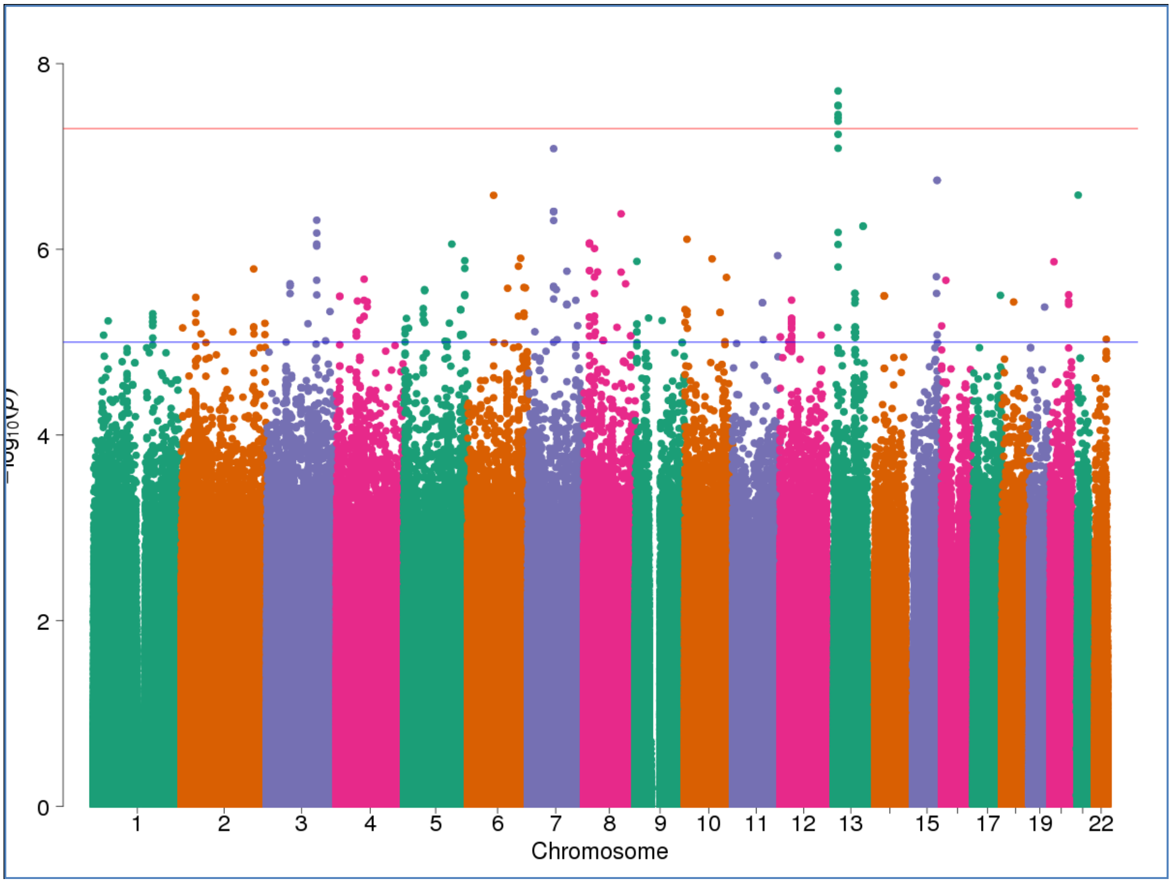

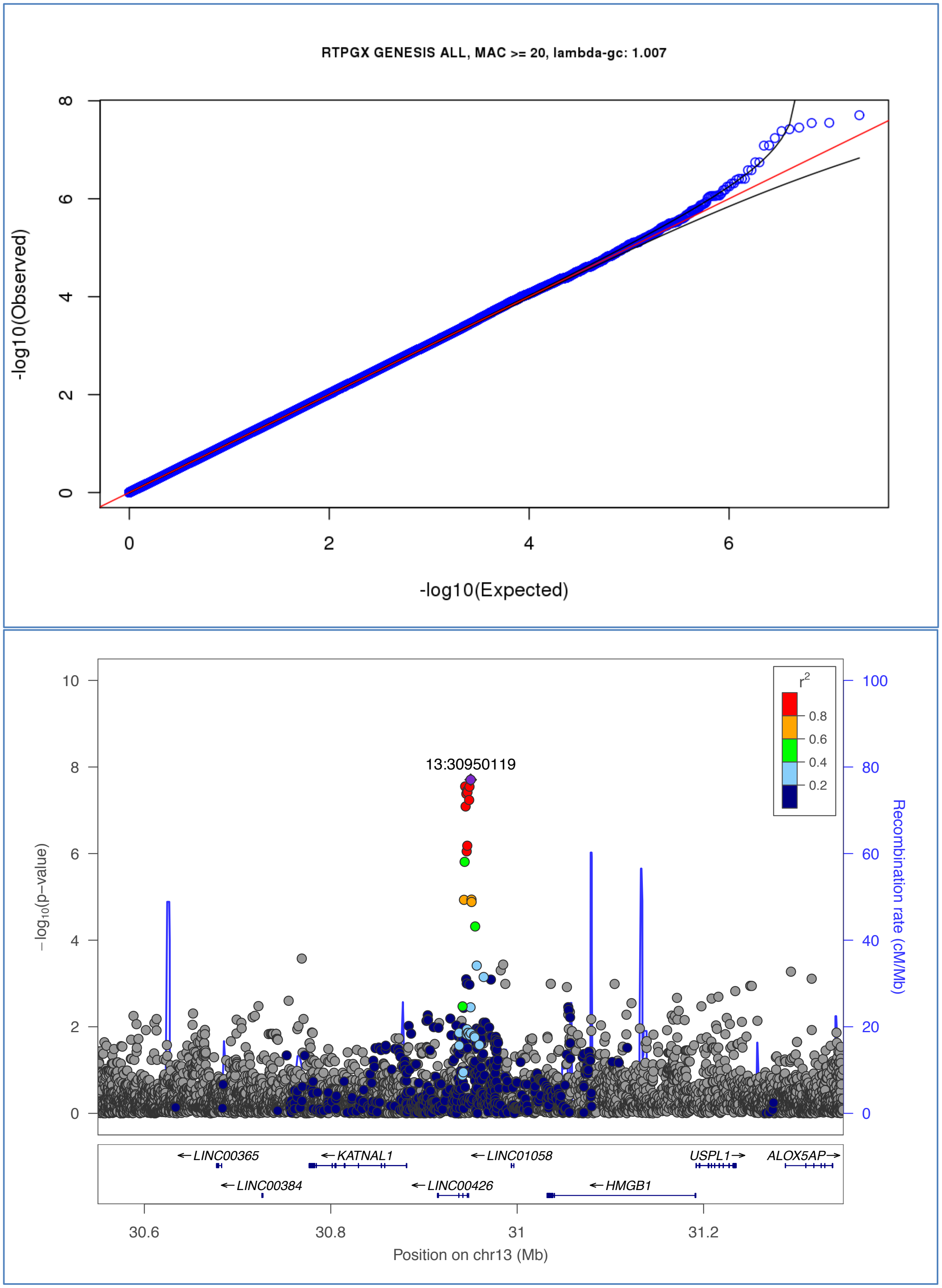


**Supplementary Figure 7**. Manhattan plot and QQ-plot of the GWAS performed with the delta MAP phenotype and the full cohort. Locus zoom of the region containing 6 genome-wide significant SNPs (rs145222507, rs1275187, rs1275192, rs1275189, rs1275191, rs2149850).

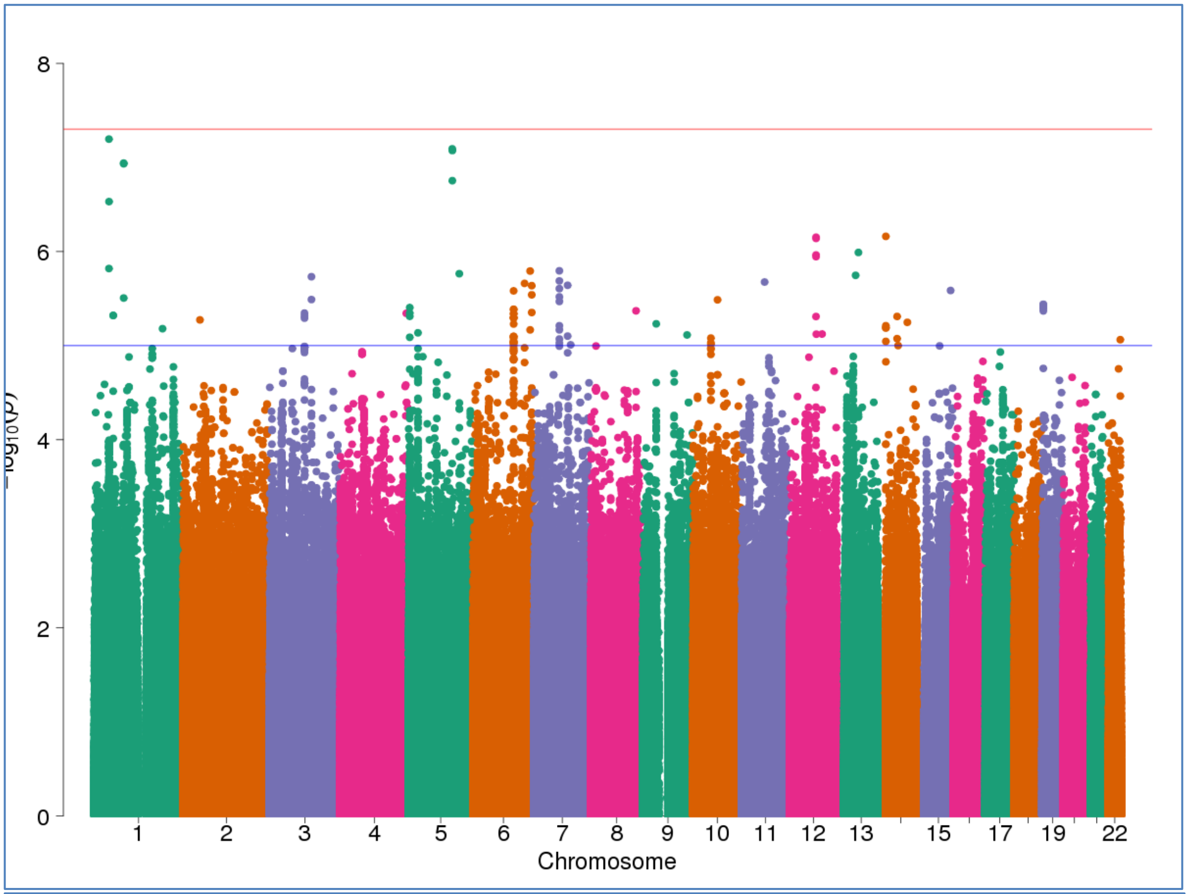


RTPGX GENESIS EA, MAC >= 20, lambda-gc: 0.999

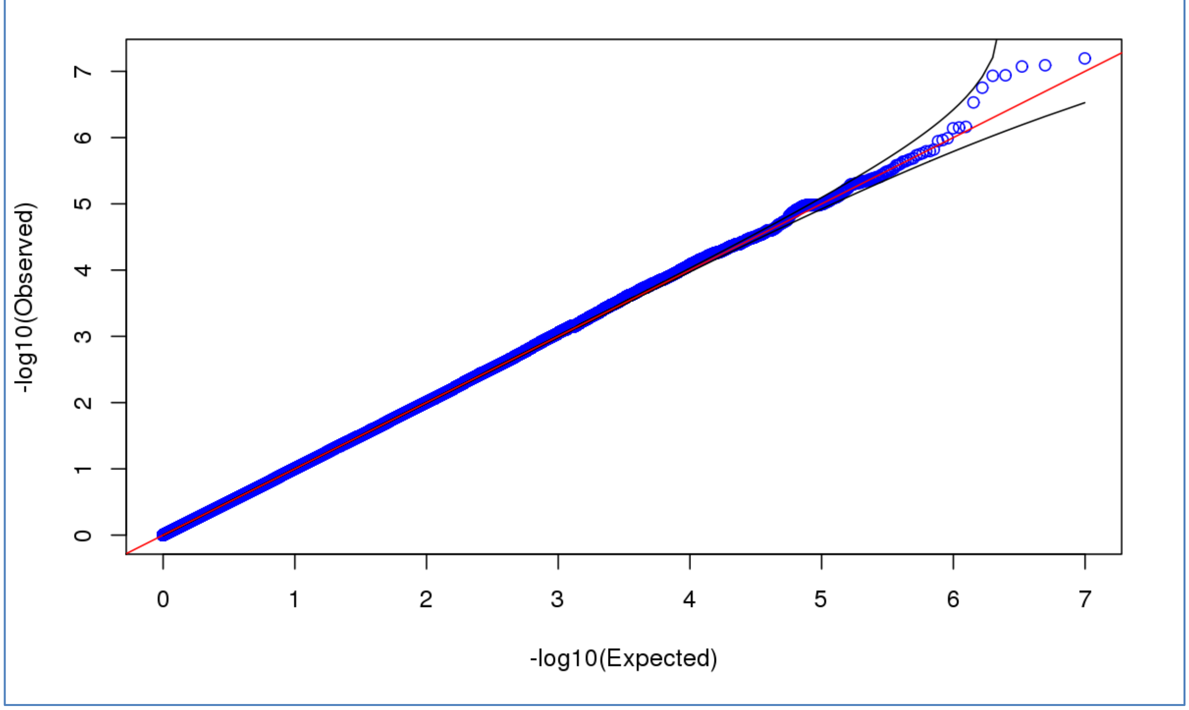

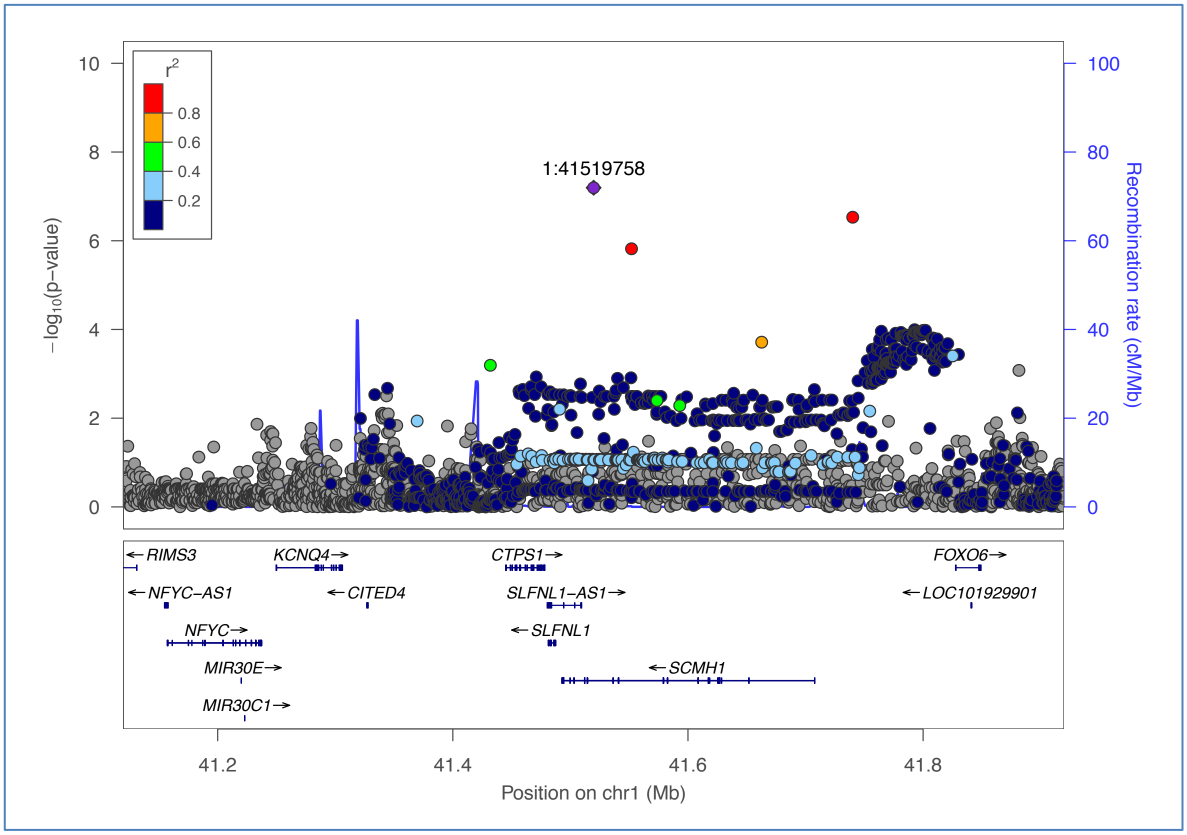


**Supplementary Figure 8**. Manhattan plot and QQ-plot of the GWAS performed with the delta MAP phenotype and the European American cohort. Locus zoom of the region containing the top SNP (rs111908123).

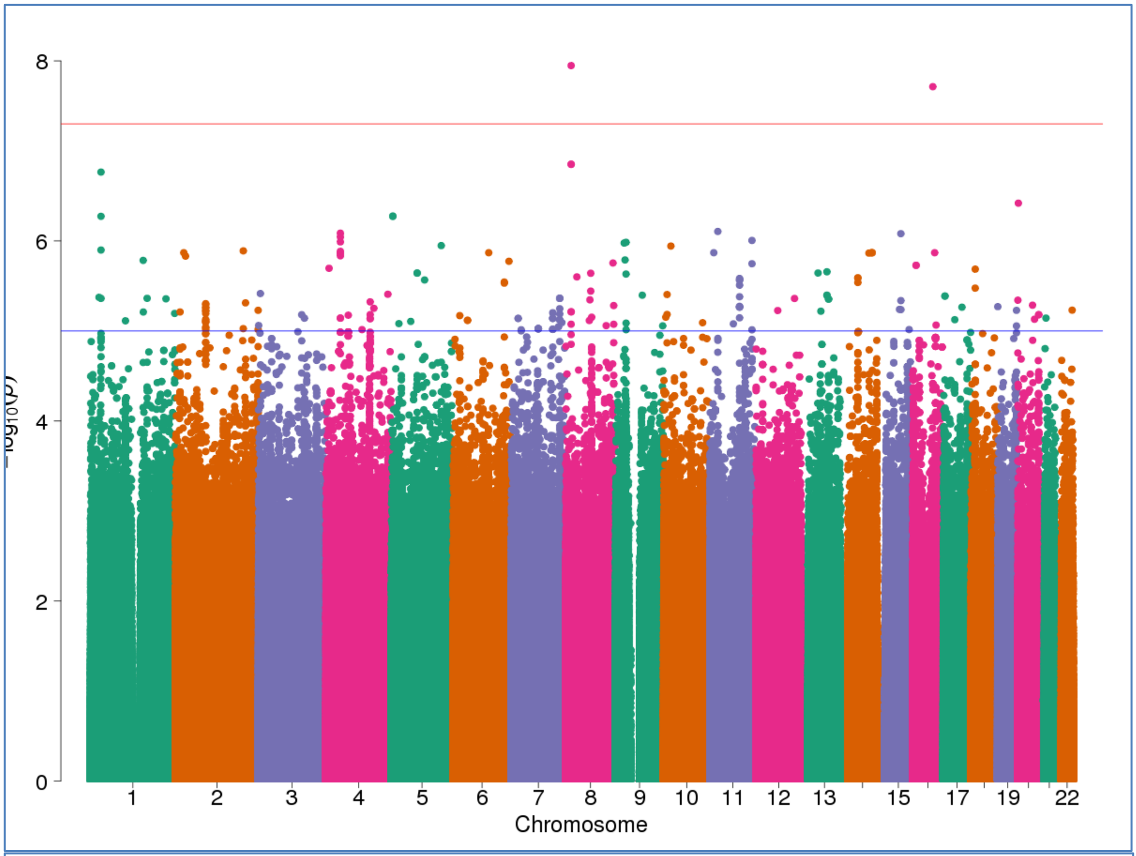

8
6
4
2
0
-log10(p)
1
2
3
4
5
6
7
8
9
10
11
12
13
15
17
19
22
Chromosome


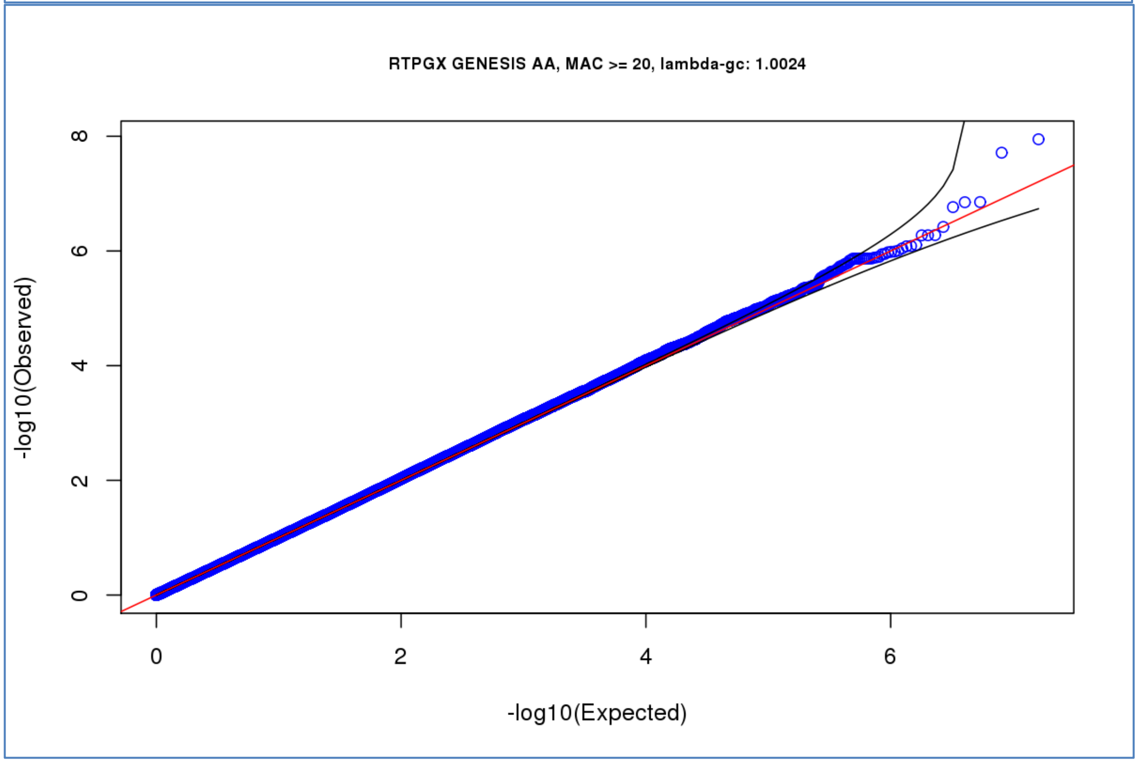

RTPGX GENESIS AA, MAC >= 20, lambda-gc: 1.0024
-log10(Observed)
0
2
4
6
8
-log10(Expected)

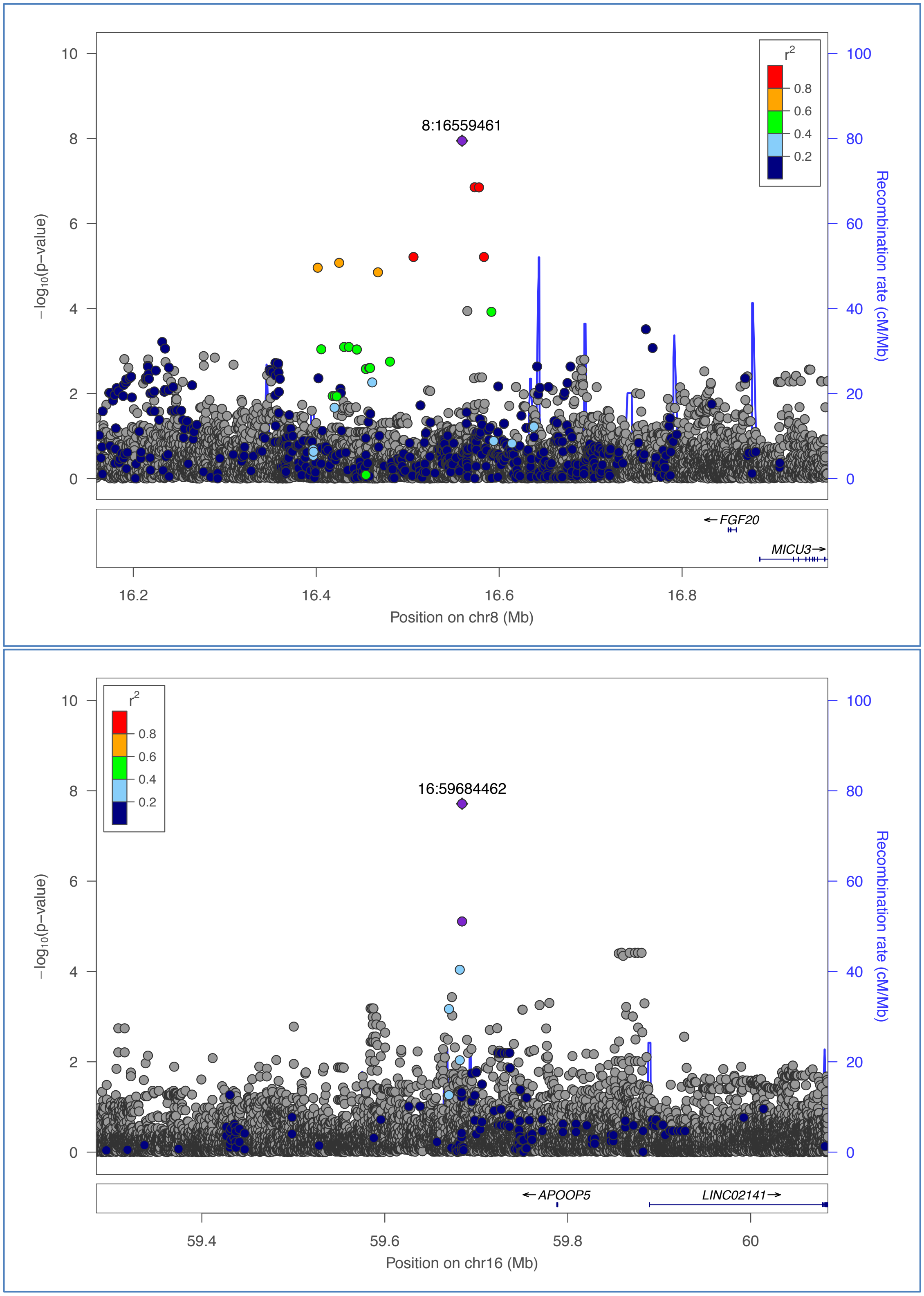


**Supplementary Figure 9**. Manhattan plot and QQ-plot of the GWAS performed with the delta DBP phenotype and the African American cohort. Locus zoom plots of the two regions containing the two genome-wide significant SNPs (rs146535276, rs143947120).

### Regeneron Genetics Center Banner Author List and Contribution Statements

All authors are listed in alphabetical order.

**RGC Management and Leadership Team**

Goncalo Abecasis, Ph.D., Aris Baras, M.D., Michael Cantor, M.D., Giovanni Coppola, M.D., Aris Economides, Ph.D., John D. Overton, Ph.D., Jeffrey G. Reid, Ph.D., Alan Shuldiner, M.D.

Contribution: All authors contributed to securing funding, study design and oversight, and review and interpretation of data and results. All authors reviewed and contributed to the final version of the manuscript.

**Sequencing and Lab Operations**

Christina Beechert, Caitlin Forsythe, M.S., Erin D. Fuller, Zhenhua Gu, M.S., Michael Lattari, Alexander Lopez, M.S., John D. Overton, Ph.D., Thomas D. Schleicher, M.S., Maria Sotiropoulos Padilla, M.S., Karina Toledo, Louis Widom, Sarah E. Wolf, M.S., Manasi Pradhan, M.S., Kia Manoochehri, Ricardo H. Ulloa.

Contribution: C.B., C.F., K.T., A.L., and J.D.O. performed and are responsible for sample genotyping. C.B, C.F., E.D.F., M.L., M.S.P., K.T., L.W., S.E.W., A.L., and J.D.O. performed and are responsible for exome sequencing. T.D.S., Z.G., A.L., and J.D.O. conceived and are responsible for laboratory automation. M.P., K.M., R.U., and J.D.O are responsible for sample tracking and the library information management system.

**Genome Informatics**

Xiaodong Bai, Ph.D., Suganthi Balasubramanian, Ph.D., Leland Barnard, Ph.D., Andrew Blumenfeld, Yating Chai, Ph.D., Gisu Eom, Lukas Habegger, Ph.D., Young Hahn, Alicia Hawes, B.S., Shareef Khalid, Jeffrey G. Reid, Ph.D., Evan K. Maxwell, Ph.D., John Penn, M.S., Jeffrey C. Staples, Ph.D., Ashish Yadav, M.S.

Contribution: X.B., A.H., Y.C., J.P., and J.G.R. performed and are responsible for analysis needed to produce exome and genotype data. G.E., Y.H., and J.G.R. provided compute infrastructure development and operational support. S.K., S.B., and J.G.R. provide variant and gene annotations and their functional interpretation of variants. E.M., L.B., J.S., A.B., A.Y., L.H., J.G.R. conceived and are responsible for creating, developing, and deploying analysis platforms and computational methods for analyzing genomic data.

**Clinical Informatics**

Nilanjana Banerjee, Ph.D., Michael Cantor, M.D.

Contribution: All authors contributed to the development and validation of clinical phenotypes used to identify study subjects and (when applicable) controls.

**Analytical Genomics and Data Science**

Goncalo Abecasis, Ph.D., Amy Damask, Ph.D., Lauren Gurski, Alexander Li, Ph.D., Nan Lin, Ph.D., Daren Liu, Jonathan Marchini Ph.D., Anthony Marcketta, Shane McCarthy, Ph.D., Colm O'Dushlaine, Ph.D., Charles Paulding, Ph.D., Claudia Schurmann, Ph.D., Dylan Sun, Tanya Teslovich, Ph.D., Cristopher Van Hout, Ph.D., Bin Ye

Contribution: Development of statistical analysis plans. QC of genotype and phenotype files and generation of analysis ready datasets. Development of statistical genetics pipelines and tools and use thereof in generation of the association results. QC, review and interpretation of result. Generation and formatting of results for manuscript figures. Contributions to the final version of the manuscript.

**Therapeutic Area Genetics**

Jan Freudenberg, M.D., Nehal Gosalia, Ph.D., Claudia Gonzaga-Jauregui, Ph.D., Julie Horowitz, Ph.D., Kavita Praveen, Ph.D.

Contribution: Development of study design and analysis plans. Development and QC of phenotype definitions. QC, review, and interpretation of association results. Contributions to the final version of the manuscript.

**Planning, Strategy, and Operations**

Paloma M. Guzzardo, Ph.D., Marcus B. Jones, Ph.D., Lyndon J. Mitnaul, Ph.D.

Contribution: All authors contributed to the management and coordination of all research activities, planning and execution. All authors managed the review of data and results for the manuscript. All authors contributed to the review process for the final version of the manuscript.

### Charles Bronfman Institute for Personalized Medicine Genomics Group Banner Author List and Contribution Statements

All authors are listed in alphabetical order.

**IPM Management and Leadership team**
Judy H. Cho, MD, Ron Do, PhD, Eimear E. Kenny, PhD, Ruth J.F. Loos, PhD, Girish Nadkarni, MD, MPH, CPH, Aniwaa Owusu Obeng, PharmD, Quingbin Song, MD, MSc.

Contribution: All authors contributed to the management and coordination of all research activities, planning and execution. All authors contributed to securing funding, study design and oversight, and review and interpretation of data and results. All authors reviewed and contributed to the final version of the manuscript.

**Data transfer and validation**
Steve Ellis, Bhupender Thakur, Quingbin Song, MD, MSc, Stephane Wenric, PhD, Lisheng Zhou, PhD

Contribution: This team coordinated the downloading of all data files, setting up of storage requirements, and verification of data fidelity to the original.

**Genotyping data quality control**
Gillian Belbin, PhD, Lisheng Zhou, PhD

Contribution: This team assessed and filtered the genome-wide array data by executing a multistep data evaluation and cleansing pipeline, identifying and removing low-quality variants and samples, and ensuring high overall quality in the resultant dataset.

**Exome data quality control**
Amanda Dobbyn, PhD, Stephane Wenric, PhD, Lisheng Zhou, PhD

Contribution: This team performed detailed investigation of quality metrics and statistics to determine what parts of the data met and diverged from expectations, including transition-transversion ratio, sequencing coverage, and Hardy-Weinberg equilibrium, and removing low-quality variants and samples accordingly.

**Samples demographics characteristics quality control**
Steve Ellis, Arden Moscati, PhD, Girish Nadkarni, MD, MPH, CPH, Rajiv Nadukuru, Stephane Wenric, PhD

Contribution: This team contributed to the identification and/or removal of study subjects with various demographics and clinical discrepancies or abnormalities. Integration of multiple sources of data allowed the identification and rectification of sample swaps, and informed what actions to take for duplicate and discrepant samples.
